# Patient-Derived Organoids Functionally Stratify Epithelial Ovarian Cancer into Clinically Relevant Chemotherapy Response Phenotypes

**DOI:** 10.64898/2026.06.10.731260

**Authors:** Shruthi Ragothaman, Raghuveer Reddy, Shelma Cheeran Sajan, Leah Achsah John, Vismaya Biju, Vipulachandra Y, Satish Sankaran, Rohit R Ranade, Smitha P.K

## Abstract

**Background:** Ovarian cancer (OC) exhibits substantial heterogeneity in response to platinum-based chemotherapy, resulting in variable clinical outcomes and frequent recurrence. Current biomarkers, including serum CA-125 kinetics and BRCA mutational status, incompletely predict therapeutic response. We investigated whether patient-derived organoids (PDOs) could functionally stratify chemotherapy sensitivity and better reflect patient-specific clinical behaviour.

**Methods:** Twenty patients with OC treated between January 2024 and May 2026 were included, from whom fourteen PDO lines were successfully established. Eight PDOs with robust low-passage expansion and comprehensive longitudinal follow-up underwent functional profiling against carboplatin, paclitaxel, olaparib, and doxorubicin. Drug responses were assessed using half-maximal inhibitory concentration (IC_50_) and area under the curve (AUC) analyses and integrated with radiological response, serum CA-125 kinetics, BRCA status, and progression-free survival (PFS).

**Results:** Clinical outcomes varied considerably despite similar platinum-taxane regimens. Although post-treatment CA-125 reduction was associated with prolonged PFS, neither CA-125 kinetics nor BRCA mutational status consistently predicted therapeutic response. PDO-guided functional stratification segregated tumours into four clinically relevant platinum-taxane response phenotypes: dual-sensitive, platinum-sensitive/taxane-resistant, platinum-resistant/taxane-sensitive, and dual-resistant. These functional categories closely mirrored radiological response, CA-125 normalisation, and disease progression patterns. PDOs exhibiting low IC_50_ and AUC values were associated with durable clinical benefit, whereas resistant PDOs tracked with persistent disease and early recurrence.

**Conclusions:** PDO-guided functional stratification captures clinically meaningful therapeutic heterogeneity in OC and complements conventional biomarkers by directly measuring tumour-specific drug susceptibility. Prospective integration of PDO testing may facilitate patient-specific therapeutic selection and support functional precision oncology approaches in ovarian cancer.

## 1. Introduction

Ovarian cancer (OC) remains the leading cause of gynecologic cancer mortality worldwide, a consequence largely driven by late-stage diagnosis and a high propensity for recurrence following cytoreductive surgery [1–3]. Although standard first-line regimens comprising platinum- and taxane-based chemotherapies initially yield high objective response rates, a substantial subset of patients inevitably develops chemoresistant disease, leading to clinical relapse and poor long-term survival [3–5]. Despite recent therapeutic advances in surgical techniques, novel targeted agents, and maintenance strategies such as poly (ADP-ribose) polymerase (PARP) inhibitors, durable clinical benefits remain limited for a significant patient cohort [6,7]. This therapeutic gap underscores the need to better understand the underlying biological drivers of drug response and to establish robust predictive frameworks.

A major challenge in contemporary OC management especially in epithelial ovarian cancer (EOC) is the marked variation in therapeutic response observed among patients receiving uniform treatment regimens [4,8,9]. While genomic biomarkers, including *BRCA1/2* mutations and homologous recombination deficiency (HRD) scores, successfully guide the selection of PARP inhibitors, their capacity to predict sensitivity or resistance to conventional cytotoxic platinum-taxane combinations remains limited [10,11]. Clinical outcomes frequently diverge from baseline genomic profiles, demonstrating that therapeutic response is driven by dynamic, functional biological processes rather than static molecular alterations [12–14]. Similarly, while serum CA-125 kinetics serve as a reliable surrogate marker for fluctuating tumor burden, they fail to provide mechanistic insights into intrinsic or acquired chemoresistance[15]. Consequently, clinical decision-making continues to rely on empirical protocols and macro-level clinicopathological features rather than an individualized, direct assessment of patient-specific tumor vulnerability.

This predictive deficit is particularly pronounced within advanced-stage clinical workflows. Whether patients are directed to neoadjuvant chemotherapy (NACT) followed by interval debulking surgery (IDS), or primary debulking surgery (PDS) followed by adjuvant chemotherapy (ACT), systemic cytotoxic therapy is almost universally initiated without prior knowledge of the underlying tumor drug-sensitivity profile [16–18]. This empirical paradigm exposes patients harboring intrinsically chemoresistant disease to ineffective toxicities while unnecessarily delaying potentially effective alternative interventions [6,19]. Across both standard surgical pathways, the absence of predictive modalities capable of assaying functional chemosensitivity prior to treatment initiation remains a critical bottleneck in oncology care.

Functional precision oncology represents a pragmatic strategy to overcome these diagnostic limitations. Patient-derived organoids (PDOs) function as *ex vivo* three-dimensional biomimetic models that preserve the histological, molecular, and phenotypic architecture of the parental malignancy, thereby maintaining both interpatient heterogeneity and differential pharmacological responses [20–22]. Crucially, unlike static genomic sequencing, PDO-based functional drug screening permits the direct, real-time quantification of viable tumor cell sensitivity to specific therapeutic agents. This approach provides a dynamic phenotypic readout of treatment efficacy, potentially exposing complex, multi-factorial resistance mechanisms that evade conventional molecular and clinical biomarkers. Emerging evidence indicates that PDOs can accurately mirror clinical outcomes across diverse solid malignancies, offering a translatable pathway toward clinically actionable, individualized therapeutic selection[21,22].

In the present study, we integrated *ex vivo* PDO functional drug testing with longitudinal clinical outcome data to analyze treatment-response heterogeneity in OC. PDO lines were successfully established from a cohort of twenty patients, from which a core subset of eight cases characterized by complete longitudinal clinical annotations underwent comprehensive functional drug profiling. By correlating *ex vivo* drug sensitivity metrics with longitudinal radiographic responses, serum CA-125 kinetics, and progression-free survival (PFS), we evaluated the utility of PDOs to functionally stratify tumors into four distinct platinum-taxane response phenotypes. Our findings demonstrate a robust concordance between *ex vivo* PDO responses and patient clinical trajectories, validating PDO-guided stratification as a powerful, complementary tool to conventional genomic biomarkers for precision oncology in OC.

## 2. Materials and methods

### 2.1. Clinical cohort and patient selection

Twenty patients diagnosed with OC were prospectively enrolled at Narayana Health City (Bengaluru, India) between January 2024 and May 2026. PDOs were successfully established from 14 cases. Downstream functional drug profiling was performed on a core subset of eight PDO lines selected based on robust, reproducible growth kinetics and the availability of comprehensive longitudinal clinical follow-up data, capturing approximately 90% of the definitive treatment outcome information required for translational matching (**Supplementary Fig. 1**).

Clinicopathological variables including age, FIGO stage, histological subtype, primary treatment sequence (PDS versus NACT), maintenance regimens, objective radiological response, and serum CA-125 kinetics were extracted from electronic medical records. PFS was defined as the interval from the final dose of chemotherapy to documented disease progression or last clinical follow-up. This study was approved by the Narayana Hrudayalaya Ethics Committee (NHH/AEC CL2024 1334), adhered strictly to the principles of the Declaration of Helsinki, and was conducted after obtaining written informed consent from all participants.

### 2.2. Tumour sample collection and PDO establishment

Fresh tumor specimens were harvested from primary ovarian or fallopian tube lesions, or from metastatic deposits within the omentum or peritoneum during PDS, IDS, or diagnostic biopsy. Tissues were processed within 2 hours of surgical excision. Samples underwent mechanical mincing followed by enzymatic dissociation with collagenase type IV to isolate epithelial-enriched cell fractions for PDO establishment, following previously established protocols [21,23]. These suspensions were embedded in growth factor-reduced extracellular matrix (ECM) and maintained in a defined, optimized ovarian cancer organoid medium under standard humidified conditions (37°C, 5% CO_2_). PDO formation and morphology were tracked via bright-field microscopy, with lines passaged every 10-14 days upon reaching confluence. Established lines were cryopreserved and subsequently thawed to validate long-term viability and protocol stability. PDO establishment efficiency was calculated as the percentage of processed specimens that yielded expandable cultures sustained beyond three consecutive passages. Detailed processing workflows, organoid media components, and cryopreservation steps are provided in the **Supplementary 1**.

### 2.3. Histological and immunophenotypic characterization

To confirm that the organoids accurately recapitulated parental tumor architecture, matched primary tissue specimens and corresponding PDOs were evaluated by hematoxylin and eosin (H&E) staining and immunohistochemistry (IHC) for diagnostic ovarian lineage markers (PAX8 and WT1) alongside the stemness-associated marker CD44. Whole-mount immunofluorescence (IF) was performed on intact organoids to characterize the expression of lineage-specific markers. Explicit staining protocols, antibody specifications, and imaging parameters are detailed in the **Supplementary 1.**

### 2.4. Functional drug profiling assays

The PDOs were dissociated into single-cell suspensions, seeded them into 96-well plates using a 1:1 matrix-to-medium ratio. Organoids were treated with carboplatin, paclitaxel, doxorubicin, and olaparib across serial dilutions ranging from 0.01 to 100 µM as monotherapies. After 120 hours of exposure, cell viability was quantified via XTT reduction assays and normalized to vehicle controls. Cytotoxicity profiles were qualitatively validated in parallel using calcein-AM and ethidium homodimer-1 fluorescence live/dead staining [21,23,24]. Full reagent sourcing, technical instrumentation specifications, and assay conditions are detailed in the **Supplementary 1.**

### 2.5. PDO-based drug response stratification

Dose-response curves were modelled using a four-parameter logistic regression algorithm to calculate the IC_50_ and AUC values for each agent. *Ex vivo* functional sensitivity to carboplatin and paclitaxel was determined by bench-marking individual PDO values against established, clinically relevant plasma concentrations (Cmax). Based on these thresholds, we stratified the PDOs into four distinct functional response phenotypes consisting of dual-sensitive, platinum-sensitive/taxane-resistant, platinum-resistant/taxane-sensitive, and dual-resistant groups. These functional classifications were subsequently integrated with patient-specific radiological data, CA-125 kinetics, PFS intervals, and baseline mutational status to determine functional-clinical translation concordance.

### 2.6. Statistical analysis

Clinical treatment responses were categorized according to RECIST version 1.1 criteria. Continuous variables and paired pre- and post-treatment CA-125 serum values were analyzed using the non-parametric Wilcoxon signed-rank test. Categorical clinical variables and correlations with mutational status were assessed using Fisher’s exact test. Two-sided -values were considered statistically significant. All statistical computations and curve-fitting modeling were executed using GraphPad Prism (v8.4.2) and Python (v3.12.7), with extended workflow parameters provided in the **Supplementary 1.**

## 3. Results

### 3.1. Establishment and characterization of Patient-Derived Organoids

PDOs were successfully established from 14 of 20 distinct OC specimens, yielding an initial derivation efficiency of 70%. All established lines demonstrated robust proliferative capacity and were sustained as expandable cultures beyond three consecutive passages (>P3). Prior to mechanical and enzymatic tissue dissociation, immunohistochemical (IHC) evaluation of formal-fixed paraffin-embedded (FFPE) source specimens confirmed high tumor cell cellularity across all 14 successful cases (**Supplementary Fig. S2**). Within 3 to 6 days of initial culture, isolated primary tumor cells in ECM droplets developed into macroscopically visible 3D organoid lines (**Supplementary Fig. S3**). These lines retained stable viability, predictable expansion kinetics, and architectural integrity across serial passages and post-cryopreservation recovery cycles. Bright-field microscopy captured pronounced inter-patient morphological heterogeneity within the cohort. Based on their distinct phenotypes, we categorized the organoids into four structural groups consisting of glandular-lumen architectures, branching/multilobulated networks, highly dense/compact spherical clusters, and heterogeneous/mixed architectural phenotypes. (**Supplementary Fig. S4**).

To validate biological fidelity, we compared the histology and immunophenotype of primary parental tumors with their corresponding PDOs. H&E and IHC analyses of tumor tissues, matched with IF profiling of the PDO lines, demonstrated precise preservation of parental tissue marker expression of the ovarian lineage markers PAX8 and WT1, alongside the stem cell marker CD44 (**Supplementary Fig. S5 and Supplementary Fig. S6**). These data confirm that the expanded *ex vivo* models faithfully maintain the definitive histological and molecular features of their parental tumors, providing a reliable platform for drug screening.

### 3.2. Conventional Clinical and Genomic Biomarkers Fail to Predict Therapeutic Heterogeneity

The 20-patient clinical cohort exhibited highly divergent longitudinal outcomes following standard empirical frontline platinum-taxane chemotherapy, spanning from complete radiological response to rapid, progressive disease (**Fig. 1**). To evaluate whether conventional biomarkers reliably predict these clinical trajectories, we analyzed a sub-cohort of eight PDO lines derived from patients with nearly complete, clinical annotation and long-term longitudinal follow-up data.

**Figure 1.**
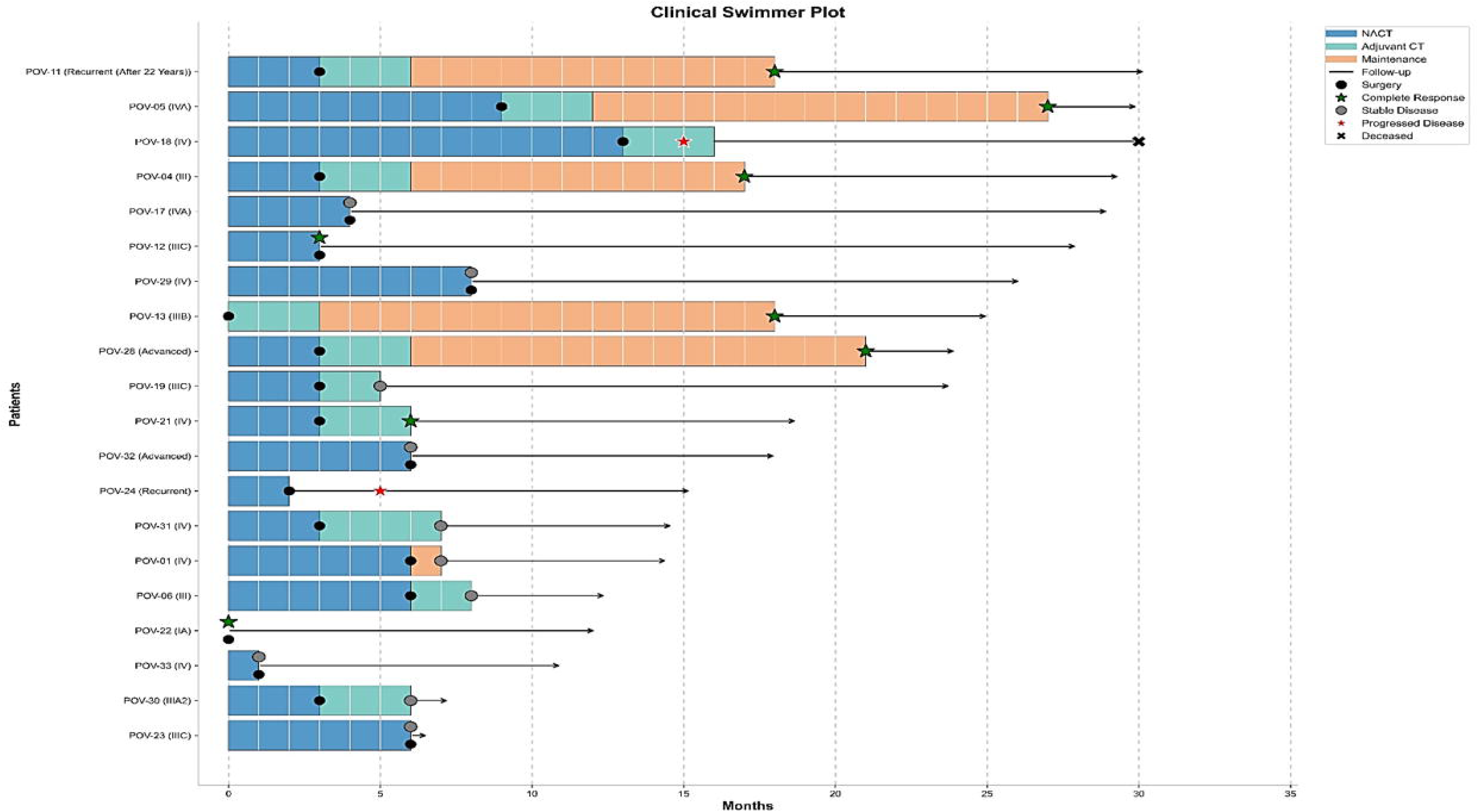
Clinical swimmer plot illustrating treatment course and outcomes of ovarian cancer patients. Swimmer plot summarising the clinical trajectories of patients with epithelial ovarian cancer over the study period. Each horizontal bar represents an individual patient and depicts the duration and sequence of treatment phases, including neoadjuvant chemotherapy (NACT; blue), adjuvant chemotherapy (teal), and maintenance therapy (orange). Black circles indicate the timing of interval or primary debulking surgery. Clinical outcomes are denoted by symbols: complete response (green star), stable disease (grey circle), progressive disease (red star), and death (black cross). Black arrows represent the duration of follow-up from diagnosis. Patients are ordered according to overall follow-up duration, illustrating the heterogeneity in treatment response, disease progression, and survival outcomes across the cohort.

Within this stratified subset, clinical responders exhibited a significant longitudinal reduction in serum CA-125 kinetics post-treatment (paired Wilcoxon signed-rank test, p = 0.0234). Conversely, patients presenting with stable or progressive disease demonstrated non-linear CA-125 fluctuations or continuous biomarker elevations[15] (**Fig. 2**).

**Figure 2.**
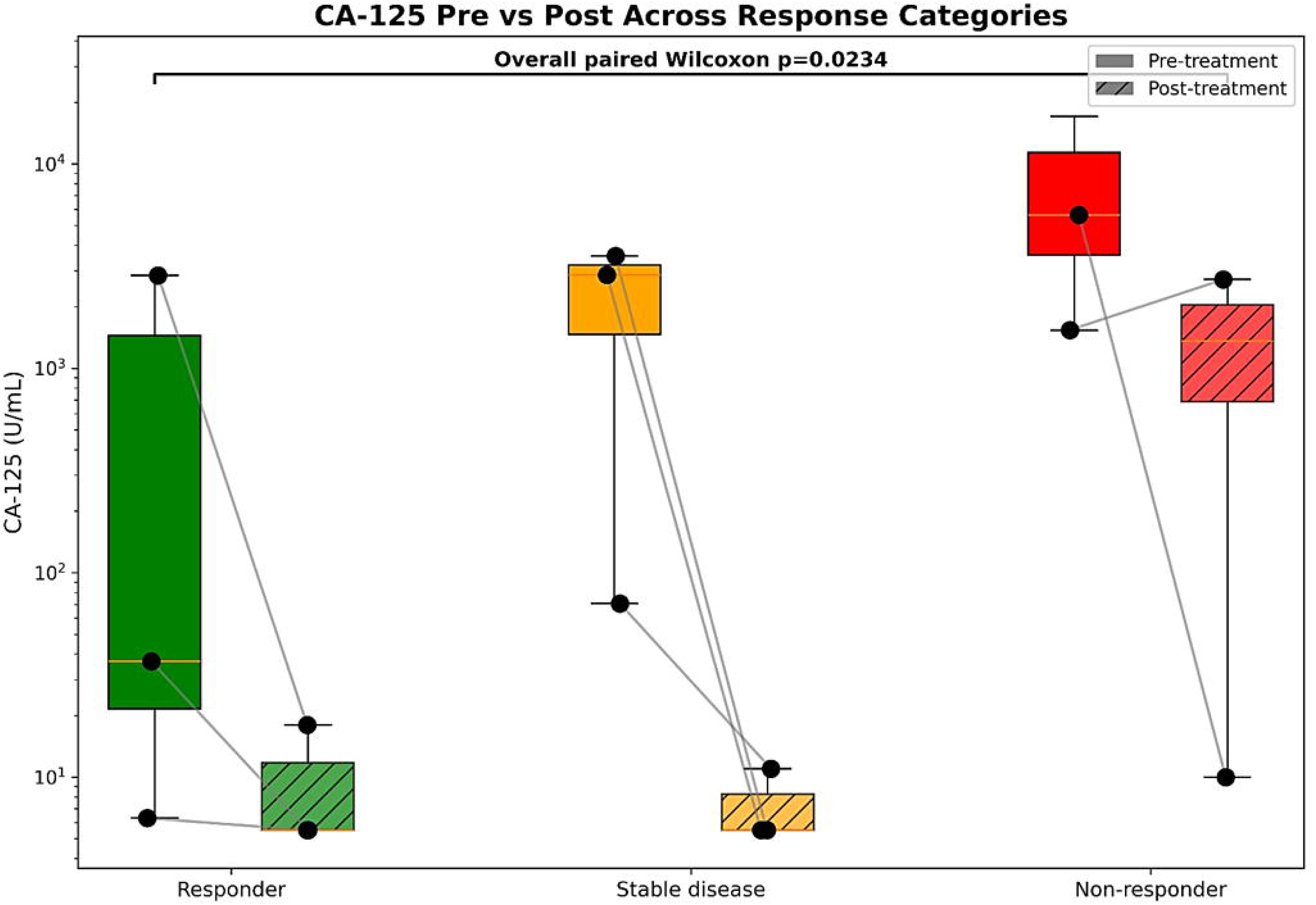
Changes in serum CA-125 levels before and after treatment according to clinical response category. Box-and-whisker plots showing pre-treatment (solid boxes) and post-treatment (hatched boxes) serum CA-125 levels (U/mL) in patients classified as responders, stable disease, and non-responders. Individual patient values are indicated by black circles and connected by grey lines to illustrate paired longitudinal changes following treatment. CA-125 values are displayed on a logarithmic scale. Responders demonstrated marked reductions in CA-125 levels following therapy, whereas patients with stable disease and non-response exhibited more variable declines and persistent elevations. Overall comparison of paired pre- and post-treatment CA-125 measurements showed a significant reduction following treatment (paired Wilcoxon signed-rank test, *p* = 0.0234).

Importantly, baseline genomic stratification failed to correlate with these clinical outcomes. Patients harboring deleterious germline *BRCA1/2* mutations, *BRCA* wild-type genotypes, and variants of uncertain significance (VUS) were distributed ubiquitously across all response tiers. No statistically significant association was observed between *BRCA* mutational status and objective clinical response (Fisher’s exact test, p = 1.0; **Supplementary Fig. S7**). These findings underscore that standard clinical markers and *BRCA* mutational status are insufficient to account for the therapeutic heterogeneity observed in patients undergoing empirical chemotherapy sequencing.

### 3.3. Ex vivo functional profiling identifies four distinct chemotherapy response phenotypes

To determine whether *ex vivo* PDO drug sensitivities recapitulate patient-specific clinical outcomes, we systematically benchmarked organoid line survival profiles against established Cmax for carboplatin (135µM) and paclitaxel (4.27µM) [25]. Dose-response profiling across the eight cohort revealed baseline variations in drug sensitivity (**Fig. 3**), categorizing the lines into four distinct functional phenotypes to establish clinical relevance.

**Figure 3.**
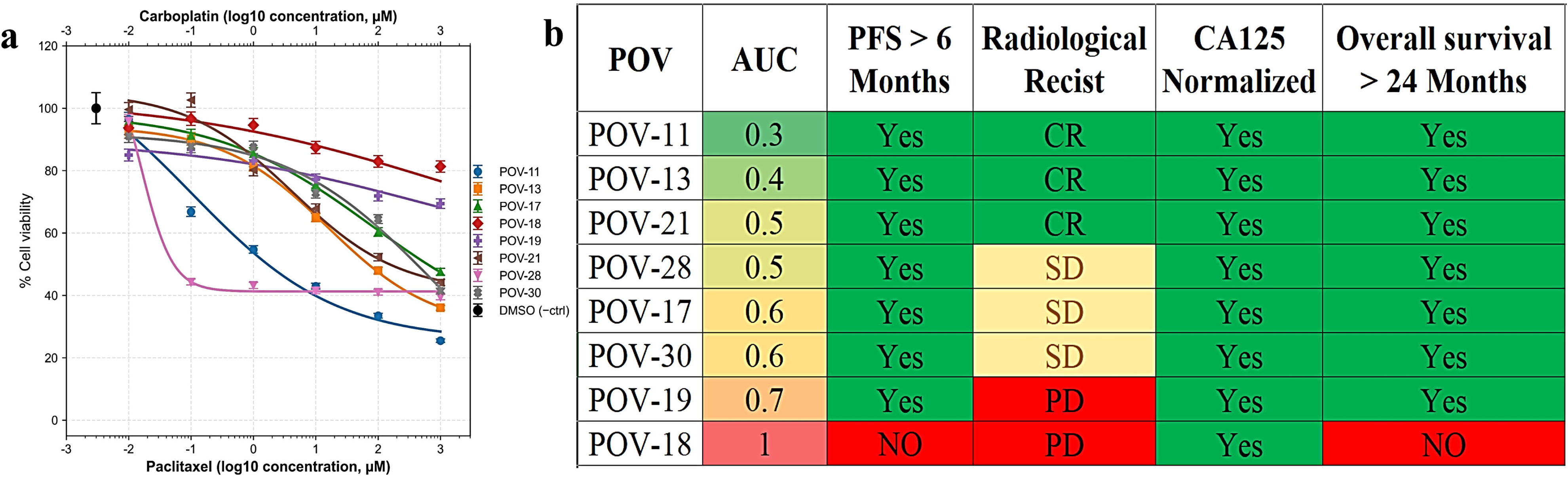
Correlation of PDO carboplatin-paclitaxel response profiles with clinical outcomes. **(a)** Dose-response curves of ovarian cancer patient-derived organoids (PDOs) treated with increasing concentrations of carboplatin-paclitaxel combinations. The lower x-axis represents paclitaxel concentrations (log_10_ µM), while the upper x-axis indicates the corresponding carboplatin concentrations (log_10_ µM). Cell viability was normalised to DMSO-treated controls and is presented as mean ± SEM. Distinct response patterns were observed across PDO lines, demonstrating inter-patient variability in sensitivity to platinum-taxane chemotherapy. **(b)** Integrated heatmap summarising PDO drug response and corresponding clinical outcomes. PDOs are ranked according to area under the curve (AUC) values, with lower AUC values indicating greater drug sensitivity. Clinical parameters include progression-free survival (PFS > 6 months), radiological response according to RECIST criteria, CA-125 normalisation following treatment, and overall survival > 24 months. Green indicates favourable clinical outcomes, yellow indicates stable disease, and red indicates adverse outcomes. Lower PDO AUC values were generally associated with improved clinical responses and survival, supporting the concordance between ex vivo PDO drug sensitivity and patient outcomes.

#### Dual platinum-taxane sensitivity

PDO models POV-11, POV-13, and POV-21 exhibited the most favourable outcomes, with sustained biochemical and radiologic responses to platinum-taxane based chemotherapy and PDO IC_50_ values were significantly below the clinical Cmax thresholds (**Fig. 4, Fig. 5 and Supplementary Fig. S8**).

**Figure 4.**
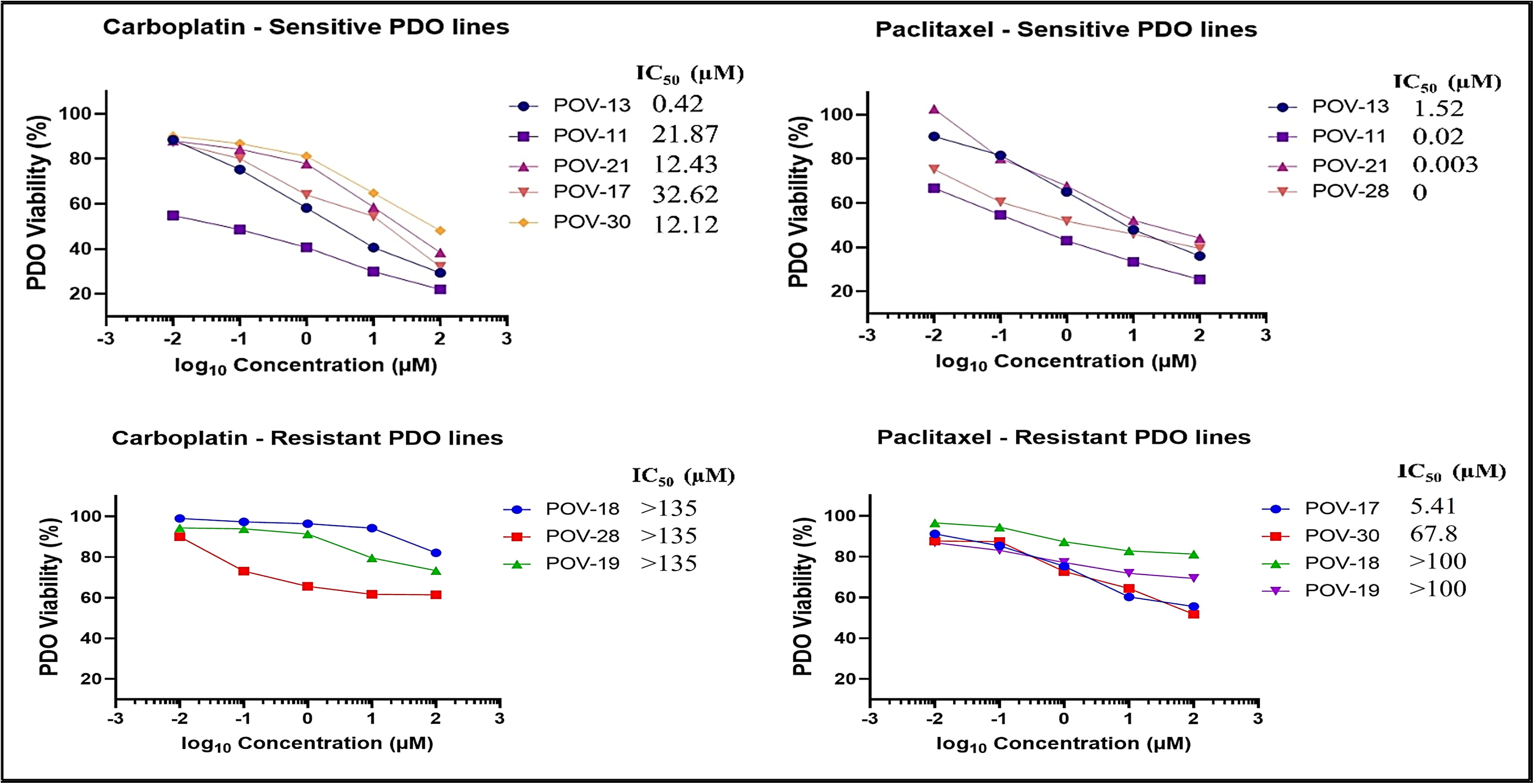
Functional stratification of ovarian cancer PDOs based on carboplatin and paclitaxel sensitivity. Dose-response curves showing viability of patient-derived organoids (PDOs) following treatment with increasing log_10_-transformed concentrations (µM) of carboplatin and paclitaxel. PDOs were classified as sensitive or resistant according to IC_50_ values relative to clinically achievable drug concentrations. Upper panels show carboplatin-sensitive (POV-13, POV-11, POV-21, POV-17, and POV-30) and paclitaxel-sensitive (POV-13, POV-11, POV-21, and POV-28) PDO lines, which exhibited a concentration-dependent reduction in viability. Lower panels show carboplatin-resistant (POV-18, POV-28, and POV-19) and paclitaxel-resistant (POV-17, POV-30, POV-18, and POV-19) PDO lines, characterised by attenuated responses across the tested concentration range. Corresponding ICCC values are indicated for each PDO line. These data demonstrate marked inter-patient variability in platinum and taxane sensitivity and identify distinct functional response phenotypes within the cohort.

**Figure 5.**
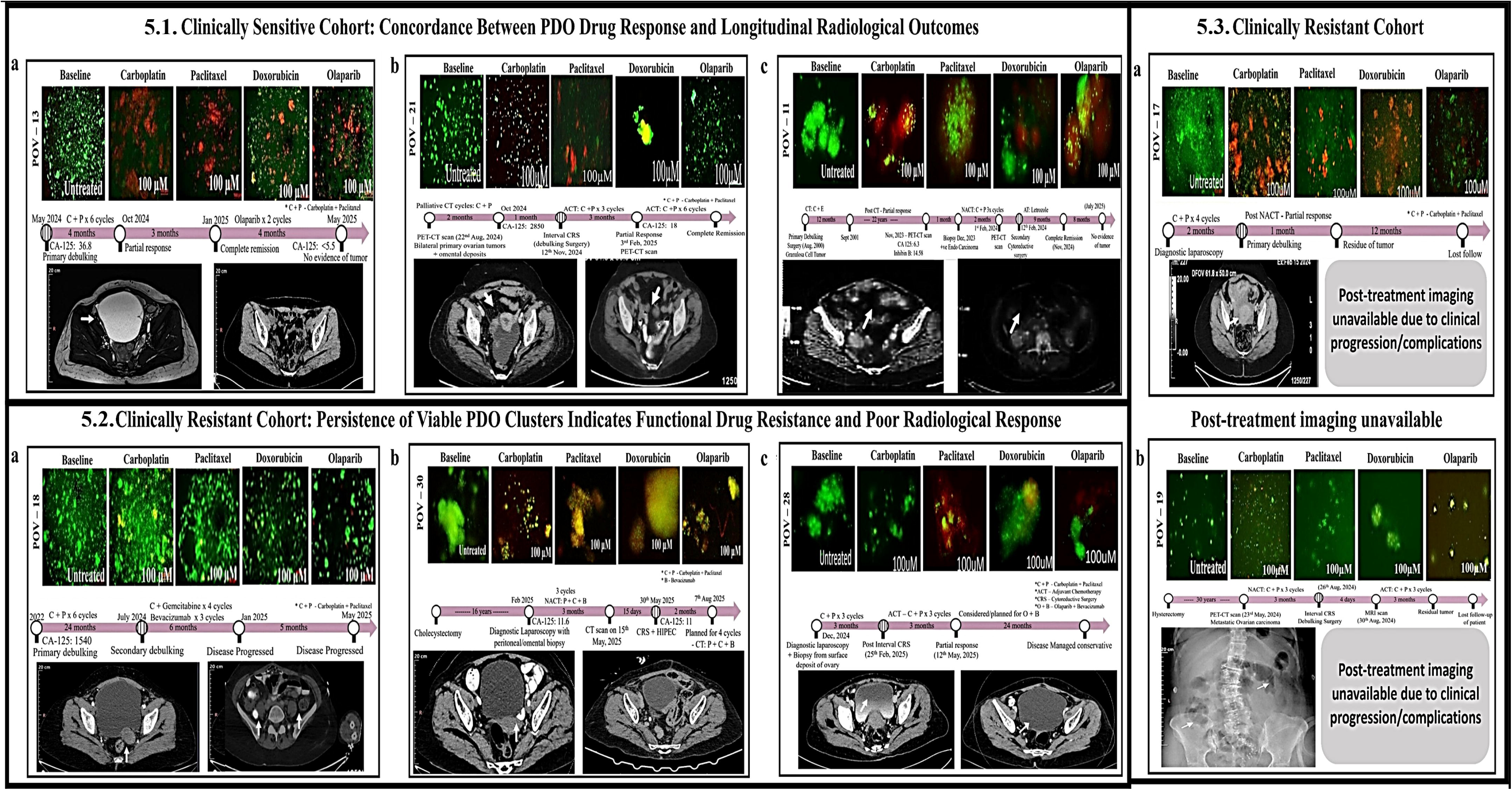
PDO drug responses recapitulate longitudinal clinical trajectories and radiological outcomes in ovarian cancer. Representative live/dead fluorescence imaging of ovarian cancer PDOs following treatment with carboplatin, paclitaxel, doxorubicin, and olaparib, integrated with corresponding clinical timelines and serial radiological assessments. Live cells are shown in green and dead cells in red. Clinical timelines summarize treatment regimens, surgical interventions, serum CA-125 dynamics, radiological responses, and disease outcomes. **(5.1) Clinically sensitive cohort.** PDOs derived from patients with favourable clinical outcomes (POV-13, POV-21, and POV-11) demonstrated marked treatment-induced cell death and disruption of organoid architecture following drug exposure. These functional responses corresponded with radiological tumour regression, CA-125 normalisation, complete or near-complete clinical responses, and prolonged disease control during follow-up. **(5.2) Clinically resistant cohort.** PDOs from patients with poor or limited clinical responses (POV-18, POV-30, and POV-28) retained viable cellular clusters despite treatment, indicating functional drug resistance. These *ex vivo* findings were concordant with persistent or recurrent disease, incomplete radiological responses, rising CA-125 levels, and disease progression observed clinically. **(5.3) Representative resistant cases with limited radiological follow-up.** PDOs from POV-17 and POV-19 exhibited persistent viability following drug exposure, consistent with treatment resistance. In these cases, post-treatment imaging was unavailable because of clinical progression, treatment-related complications, or loss to follow-up; therefore, clinical correlation was based on available treatment history and disease course. Collectively, these data demonstrate concordance between PDO-derived functional drug responses and longitudinal patient outcomes, supporting the utility of PDOs as ex vivo models for predicting therapeutic sensitivity and resistance in ovarian cancer.

POV13, a 42-year-old patient with germline BRCA1 mutated stage IIIB High-Grade Serous Ovarian Carcinoma (HGSOC), achieved complete remission following PDS and ACT before initiating olaparib maintenance. Clinical remission and clear radiological imaging directly mirrored PDO functional sensitivity to carboplatin (0.42 µM) and paclitaxel (1.52 µM), matching expected HRD phenotypes (**Fig. 5.1a**). Similarly, POV21, a 75-year-old patient with stage IV HGSOC harboring a BRCA2 variant of uncertain significance (VUS), presented with widespread metastatic disease. Following NACT and IDS the patient achieved complete remission with no evidence of disease on PET-CT imaging at follow-up (**Fig. 5.1b**). This profound clinical response aligned with matched PDO assays demonstrating pronounced *ex vivo* sensitivity to carboplatin (12.43 µM) and paclitaxel (0.003 µM).

Conversely, POV11, a 64-year-old patient with recurrent stage IA endometrioid ovarian carcinoma, presented after an exceptional 22-year platinum free interval (PFI). Despite the absence of canonical homologous recombination deficiencies (HRD) and detectable BRCA1/2 alterations, following IDS and platinum-based chemotherapy, the patient achieved a durable clinical remission with no radiological evidence of disease (**Fig. 5.1c**). PDO assays demonstrated matching sharp organoid sensitivity to carboplatin (IC_50_ = 21.87µM) and paclitaxel (IC_50_ = 0.02µM). These results indicate that dual chemotherapeutic sensitivity can occur across diverse molecular backgrounds independent of baseline HRD status.

#### Platinum sensitive-taxane resistant group

PDO lines POV-17 and POV-30 retained sensitivity to carboplatin but exhibited paclitaxel IC_50_ values that exceeded clinically Cmax.

POV17, a 55-year-old patient with germline BRCA1/2 wild type stage IVA HGSOC, demonstrated a partial radiologic response following NACT before being lost to follow up. The matched PDO line demonstrated moderate carboplatin sensitivity (IC_50_ = 32.62µM) but reduced susceptibility to paclitaxel (IC_50_ = 5.41µM), crossing the Cmax threshold indicating functional taxane resistance (**Fig. 5.3a**). Similarly, patient POV30, a 75-year-old with HRP stage IIIA2 HGSOC, received NACT combined with bevacizumab, followed by IDS and hyperthermic intraperitoneal chemotherapy. Post treatment imaging confirmed Stable disease rather than complete remission. The corresponding PDOs remained responsive to carboplatin (IC_50_ = 12.12µM) but displayed marked paclitaxel resistance (IC_50_ = 67.8µM), directly capturing the constrained efficacy observed in the clinic. Live/dead fluorescence imaging corroborated these data, showing carboplatin induced organoid fragmentation and mixed fluorescence alongside persistent, highly viable cellular clusters following paclitaxel exposure (**Fig. 5.2b**). These findings highlight how intrinsic or acquired taxane resistance can limit clinical response durability despite retained platinum sensitivity.

#### Platinum resistant-taxane sensitive group

Functional tracking of line POV-28 identified an inverse, uncoupled phenotype characterized by definitive platinum resistance paired with high taxane sensitivity.

POV28, a 53-year-old patient with BRCA1 mutated, advanced low-grade serous ovarian carcinoma (LGSOC) achieved stable disease for 9 months under PARP inhibitor therapy and remains stable at the latest follow-up. Matched PDOs exhibited marked carboplatin resistance (IC_50_ > 100 µM) together with robust paclitaxel susceptibility (IC_50_ = 0.005μM) and extensive taxane-induced organoid fragmentation. This drug-response pattern aligns with the recognized therapeutic profile of LGSOC[26], a histological subtype associated with reduced platinum sensitivity and heterogeneous responses to conventional cytotoxic chemotherapy (**Fig. 5.2c**).

#### Dual platinum-taxane resistance

PDO models POV-18 and POV-19 exhibited profound cross-resistance to both frontline therapeutics, with IC_50_ values vastly exceeding established clinical Cmax concentrations. Patient POV18 (43-years-old, germline BRCA1 mutated stage IV HGSOC) experienced disease recurrence in 2024 after a 1.5-year DFI following PDS. Recurrent tumor tissue was utilized for PDO generation prior to secondary CRS, following four cycles of gemcitabine-carboplatin and three cycles of bevacizumab. The patient subsequently defaulted on ACT, experienced disease progression within six months, received a single cycle of liposomal doxorubicin-carboplatin, declined further therapy, and succumbed to the disease in April 2026. Consistently, matched PDO assays demonstrated profound resistance to carboplatin (IC_50_ > 135µM), paclitaxel (IC_50_ > 100 µM), and doxorubicin (IC_50_ > 100 µM) (**Fig. 5.2a**). Patient POV19 (68-years-old, HRP stage IIIC HGSOC) received NACT and IDS, which resulted in stable residual disease before the patient was lost to follow up at 2.5 months. Matched PDO lines yielded IC_50_ values >100 µM for both frontline agents (**Fig. 5.3b**). Live/dead fluorescence imaging substantiated these findings, showing intact, viable organoid structures with minimal fragmentation following drug exposure (**Fig. 5.2a and Fig. 6.3b, top panels**). Live/dead fluorescence imaging validated these findings, demonstrating the persistence of viable, Calcein-AM-positive organoid clusters following paclitaxel exposure, which contrasted with complete structural fragmentation in carboplatin-treated conditions (**Fig. 5**). Individual longitudinal histories are detailed in **Supplementary 2 (Table 1)**.

### 3.4. Functional Responses to Olaparib and Doxorubicin

Olaparib screening of BRCA-mutated PDOs[27] revealed profound functional heterogeneity that frequently diverged from clinical reality. Case POV-13 (germline BRCA1-mutated) exhibited moderate *in vitro* resistance (IC_50_ = 52.01 µM) while POV-28 (BRCA-mutated) displayed absolute functional resistance (IC_50_ >100 µM). Both profiles exceeded the established clinical Cmax (∼20 µM)[28]. Yet, despite these resistant in vitro thresholds, both patients achieved durable, complete clinical responses under frontline PARP inhibitor maintenance[29]. Conversely, POV-18 demonstrated concordant, profound in vitro resistance (IC_50_ >100 µM) that accurately mirrored an aggressive, recurrent clinical course. POV-17 (germline BRCA-mutated) demonstrated an intermediate phenotype, achieving stable clinical disease before the patient was lost to follow-up (**Fig. 5.2a**).

This discordance underscores both the strengths and current limitations of epithelial-enriched PDO models. Although PDOs faithfully capture tumour-intrinsic drug responses, clinical benefit from PARP inhibition may additionally be shaped by microenvironmental, immune, and pharmacological factors that are not represented *ex vivo*[30,31]. Importantly, these findings provide a rationale for developing next-generation PDO models that incorporate microenvironmental and immune complexity to better capture clinical therapeutic responses. These findings indicate that BRCA mutation status alone does not uniformly predict functional sensitivity to PARP inhibition and underscore the complexity of treatment response in OC[32,33].

To determine whether an alternative cytotoxic mechanism could overcome frontline resistance, epithelial PDOs were exposed to doxorubicin hydrochloride, which induces DNA damage through Topoisomerase II inhibition, DNA intercalation, and reactive oxygen species generation. Despite this distinct mechanism of action, PDOs exhibited profound resistance, with IC_50_ values exceeding 100μM across all resistant models (**Fig. 5.2a; Supplementary Fig. S***9*). As this phenotype was maintained in epithelial-enriched PDO cultures lacking stromal support, it suggests a predominantly tumour-intrinsic multidrug-resistant state[34]. Notably, the observed IC_50_ values far exceeded clinically Cmax of doxorubicin (1 - 5μM), indicating that conventional anthracycline therapy is unlikely to provide meaningful benefit in this frontline-resistant cohort[35].

Collectively, these findings demonstrate that PDO based functional testing stratifies tumors into clinically meaningful response groups that closely reflect real-world patient outcomes (**Supplementary Fig. S10**). Importantly, functional sensitivity patterns capture therapeutic responsiveness across diverse molecular backgrounds including BRCA1 mutated, BRCA2 VUS, and HRP-tumors providing functional insights that complement genomic markers for precision oncology.

## 4. Discussion

This study demonstrates that PDOs provide a robust functional platform for capturing the marked interpatient heterogeneity in therapeutic response observed in OC. Integration of *ex vivo* drug sensitivity data with longitudinal clinical outcomes revealed strong concordance between PDO-derived response profiles[21,22,36] and patient treatment trajectories, enabling stratification into four clinically relevant response categories: platinum-taxane sensitive, platinum-sensitive/taxane-resistant, platinum-resistant/taxane-sensitive, and platinum-taxane resistant.

Patients classified as platinum-taxane sensitive exhibited durable disease control, PFS, and favourable clinical outcomes. In contrast, platinum-taxane resistant PDOs were associated with early disease progression, recurrence, and poor clinical outcomes. Importantly, a substantial proportion of PDOs demonstrated discordant response patterns, where sensitivity to one agent coexisted with resistance to the other. These intermediate response phenotypes highlight a critical limitation of empirical platinum-taxane combination chemotherapy[6,37,38], whereby ineffective agents may contribute little therapeutic benefit while increasing treatment-related toxicity. Such findings are particularly relevant in the neoadjuvant setting, where treatment decisions are made without direct evidence of tumour-specific drug susceptibility. Exposure of intrinsically resistant tumours to standard chemotherapy may not only reduce treatment efficacy but also promote the expansion of resistant cellular populations.

While CA-125 kinetics reflected changes in tumour burden during treatment and BRCA status provided insight into DNA repair deficiencies, neither parameter consistently predicted therapeutic response across the cohort. Notably, PDO-derived drug sensitivity profiles correlated more closely with clinical outcomes than individual genomic or clinical biomarkers alone. Several cases demonstrated functional responses that were not predicted by BRCA or HRD status, emphasizing the limitations of relying solely on molecular markers in a biologically heterogeneous disease[32,39].

Beyond frontline therapy, PDO profiling identified evidence of multidrug resistance[40]. Platinum-taxane resistant organoids frequently exhibited cross-resistance to olaparib and doxorubicin despite their distinct mechanisms of action[27,29]. Doxorubicin responses were particularly notable, with IC_50_ values frequently exceeding clinically achievable plasma concentrations, suggesting limited utility of anthracycline-based therapy in this resistant subgroup. Similarly, sensitivity to olaparib was observed across diverse genomic backgrounds and did not consistently correlate with BRCA mutational status, supporting previous observations that functional drug response cannot always be inferred from genomic alterations alone.

The observation that these response phenotypes were maintained in epithelial-enriched PDO cultures, independent of stromal or immune cell contributions, suggests that resistance mechanisms are largely retained within the tumour compartment itself. Such intrinsic multidrug-resistant states may explain why some patients experience poor outcomes despite multiple lines of therapy and underscore the need for alternative therapeutic strategies beyond conventional treatment escalation.

Several limitations should be acknowledged. The relatively small cohort size limits statistical power and generalizability. Although strong concordance between PDO responses and clinical outcomes was observed, prospective validation in larger patient cohorts will be necessary to establish clinical utility. Furthermore, while PDOs preserve key biological characteristics of the original tumour, they do not fully recapitulate the complexity of the in vivo tumour microenvironment and therefore cannot capture all determinants of treatment response.

In conclusion, PDO-based functional profiling provides a clinically relevant framework for understanding therapeutic heterogeneity and identifying multidrug-resistant disease in OC. Importantly, classification of tumours into platinum-taxane sensitive, platinum-sensitive/taxane-resistant, platinum-resistant/taxane-sensitive, and platinum-taxane resistant phenotypes offer a practical approach for patient stratification and treatment optimization. Integration of PDO testing using tumour tissue obtained during diagnostic laparoscopy could enable prospective assessment of patient-specific therapeutic vulnerabilities before initiation of systemic therapy. Such an approach may facilitate more rational treatment selection, avoid ineffective therapies, and support prioritization of targeted therapies or clinical trial-directed interventions for patients unlikely to benefit from standard empirical chemotherapy. Collectively, these findings support the incorporation of functional precision oncology approaches into the clinical management of OC.

## Supporting information

Supplementary Results

## Declaration of generative AI and AI-assisted technologies in the manuscript preparation process

During the preparation of this manuscript, the authors used ChatGPT (OpenAI) to support drafting and rephrasing for clarity of sections of the text. After using this tool, the authors reviewed and edited the content as needed and take full responsibility for the content of the study.

## CRediT authorship contribution statement

Conceptualization: Smitha P.K, Rohit R. Ranade

Data curation: Shruthi Ragothaman, Smitha P.K

Formal analysis: Shruthi Ragothaman, Raghuveer Reddy, Vipulachandra Y, Smitha P.K,

Investigation: Shruthi Ragothaman, Raghuveer Reddy, Shelma Cheeran Sajan, Leah Achsah John, Vismaya Biju, Vipulachandra Y, Rohit R. Ranade Smitha P.K

Methodology: Shruthi Ragothaman, Smitha P.K

Supervision: Smitha P.K, Rohit R. Ranade, Satish Sankaran

Writing: original draft: Shruthi Ragothaman

Writing: review & editing: Smitha P.K, Satish Sankaran, Vipulachandra Y, Rohit R. Ranade

Clinical Data Collection: Vipulachandra Y

## Funding

This research was supported by institutional seed funding from Mazumdar Shaw Medical Foundation (MSMF). No external funding was received.

## Declaration of competing interest

The authors declare that the research was conducted in the absence of any commercial or financial relationships that could be construed as a potential conflict of interest.

## Data availability

The authors confirm that the data supporting the findings of this study are available within the article and its supplementary materials.

---

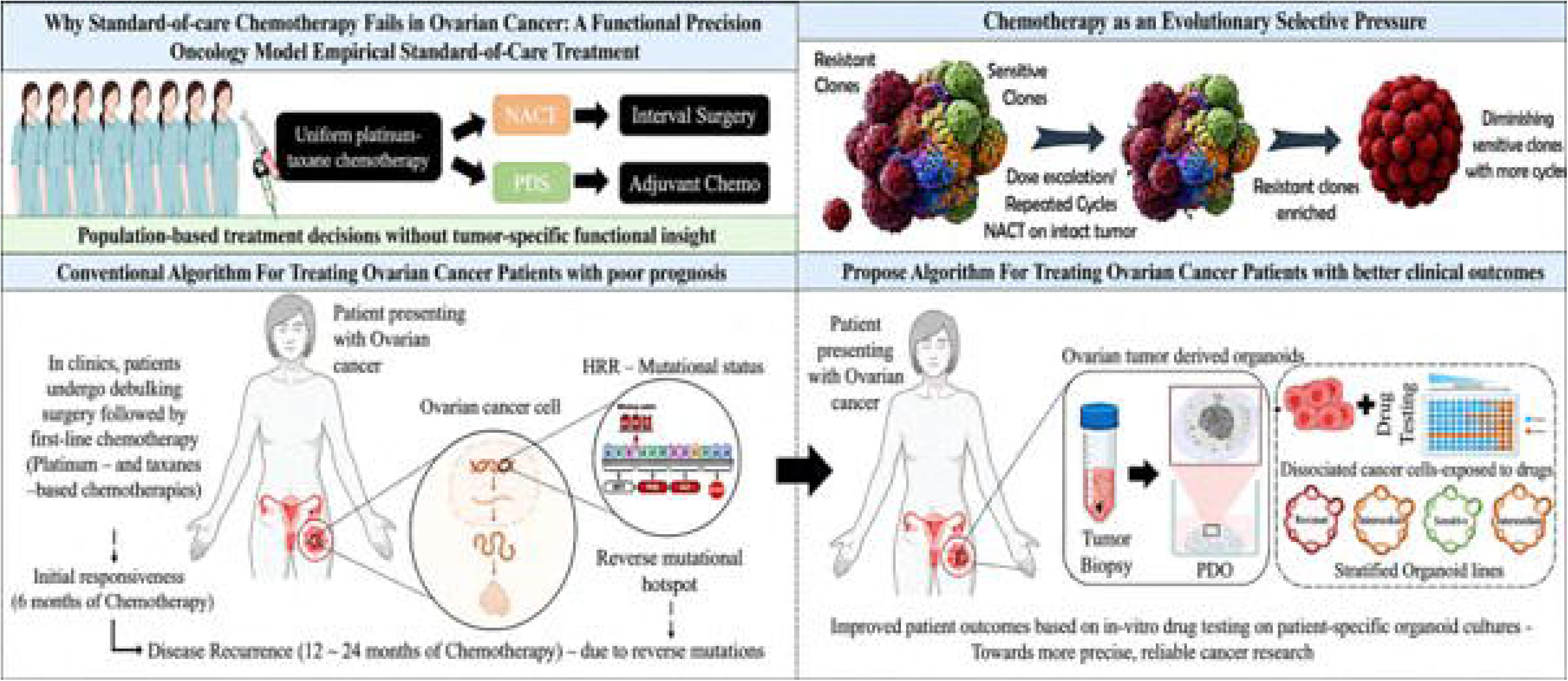

## Notes

### Competing Interest Statement

The authors have declared no competing interest.

### Summary of Updates

Format has been revised according to the submitted journal

## References

[1] Arcieri M, Tius V, Filippin S, Aletti G, Lorusso D, Fagotti A, et al. Management of Patients with Epithelial Ovarian Cancer: A Systematic Comparison of International Guidelines from Scientific Societies (AIOM-BGCS-ESGO-ESMO-JGSO-NCCN-NICE). Cancers (Basel) 2025;17:3915. 10.3390/cancers17243915.

[2] Sambasivan S. Epithelial ovarian cancer: Review article. Cancer Treat Res Commun 2022;33:100629. 10.1016/j.ctarc.2022.100629.

[3] Webb PM, Jordan SJ. Global epidemiology of epithelial ovarian cancer. Nat Rev Clin Oncol 2024;21:389–400. 10.1038/s41571-024-00881-3.

[4] Yang L, Xie H-J, Li Y-Y, Wang X, Liu X-X, Mai J. Molecular mechanisms of platinum-based chemotherapy resistance in ovarian cancer (Review). Oncol Rep 2022;47:82. 10.3892/or.2022.8293.

[5] Ren Y, Xu R, Wang Y, Su L, Su J. Global, regional, and national burden of ovarian cancer in women aged 45+from 1990 to 2021 and projections for 2050: a systematic analysis based on the 2021 global burden of disease study. J Cancer Res Clin Oncol 2025;151:225. 10.1007/s00432-025-06277-9.

[6] Wang G, Yang H, Wang Y, Qin J. Ovarian cancer targeted therapy: current landscape and future challenges. Front Oncol 2025;15. 10.3389/fonc.2025.1535235.

[7] Apelian S, Martincuks A, Whittum M, Yasukawa M, Nguy L, Mathyk B, et al. PARP Inhibitors in Ovarian Cancer: Resistance Mechanisms, Clinical Evidence, and Evolving Strategies. Biomedicines 2025;13:1126. 10.3390/biomedicines13051126.

[8] Masoodi T, Siraj S, Siraj AK, Azam S, Qadri Z, Parvathareddy SK, et al. Genetic heterogeneity and evolutionary history of high-grade ovarian carcinoma and matched distant metastases. Br J Cancer 2020;122:1219–30. 10.1038/s41416-020-0763-4.

[9] McGranahan N, Swanton C. Clonal Heterogeneity and Tumor Evolution: Past, Present, and the Future. Cell 2017;168:613–28. 10.1016/j.cell.2017.01.018.

[10] Marconato N, Tommasi O De, Paladin D, Boscarino D, Spagnol G, Saccardi C, et al. Unraveling homologous recombination deficiency in ovarian cancer: A review of currently available testing platforms. Cancers (Basel) 2025;17:1771.

[11] Witjes VM, de Hullu JA, van Remortele A, Vreede L, Rosenberg EH, Cillessen SAGM, et al. Evaluation of homologous recombination testing in ovarian carcinoma. Virchows Archiv 2026. 10.1007/s00428-026-04432-2.

[12] Cheng F, Shao F, Tian Y, Chen S. Genomic and clinical insights into ovarian cancer: subtype-specific alterations and predictors of metastasis and relapse. Discover Oncology 2025;16:907.

[13] Dong J, Ni J, Chen J, Wang X, Ye L, Xu X, et al. Genomic alteration discordance in the paired primary-recurrent ovarian cancers: based on the comprehensive genomic profiling (CGP) analysis. J Ovarian Res 2024;17:133.

[14] Diaz M, Gull N, Peng P-C, Dabke K, Baker J, Sun J, et al. The mutation and clonality profile of genomically unstable high grade serous ovarian cancer is established early in tumor development and conserved throughout therapy resistance 2025. 10.1101/2025.02.14.638365.

[15] Jitmana K, Griffiths JI, Fereday S, DeFazio A, Bowtell D, Adler FR. Mathematical modeling of the evolution of resistance and aggressiveness of high-grade serous ovarian cancer from patient CA-125 time series. PLoS Comput Biol 2024;20:e1012073. 10.1371/journal.pcbi.1012073.

[16] Gaillard S, Lacchetti C, Armstrong DK, Cliby WA, Edelson MI, Garcia AA, et al. Neoadjuvant Chemotherapy for Newly Diagnosed, Advanced Ovarian Cancer: ASCO Guideline Update. Journal of Clinical Oncology 2025;43:868–91. 10.1200/JCO-24-02589.

[17] Uçkan HH, Salman MC, Yüce K. Effect of neoadjuvant chemotherapy on primary and metastatic tumour burden in epithelial ovarian cancer: a retrospective analysis of oncologic surgeries at a university hospital. Italian Journal of Gynaecology and Obstetrics 2026. 10.36129/jog.2026.265.

[18] Abel MK, Mazina V, Bregar AJ, Gockley AA, del Carmen MG, Eisenhauer EL, et al. Neoadjuvant Chemotherapy, Case Volume, and Mortality in Advanced Ovarian Cancer. JAMA Netw Open 2025;8:e2523434. 10.1001/jamanetworkopen.2025.23434.

[19] Williams MJ, Vázquez-García I, Tam G, Wu M, Varice N, Havasov E, et al. Tracking clonal evolution during treatment in ovarian cancer using cell-free DNA. Nature 2025;647:757–65. 10.1038/s41586-025-09580-0.

[20] Wensink GE, Elias SG, Mullenders J, Koopman M, Boj SF, Kranenburg OW, et al. Patient-derived organoids as a predictive biomarker for treatment response in cancer patients. NPJ Precis Oncol 2021;5. 10.1038/s41698-021-00168-1.

[21] Kopper O, de Witte CJ, Lõhmussaar K, Valle-Inclan JE, Hami N, Kester L, et al. An organoid platform for ovarian cancer captures intra- and interpatient heterogeneity. Nat Med 2019;25:838–49. 10.1038/s41591-019-0422-6.

[22] de Witte CJ, Espejo Valle-Inclan J, Hami N, Lõhmussaar K, Kopper O, Vreuls CPH, et al. Patient-Derived Ovarian Cancer Organoids Mimic Clinical Response and Exhibit Heterogeneous Inter- and Intrapatient Drug Responses. Cell Rep 2020;31. 10.1016/j.celrep.2020.107762.

[23] Maenhoudt N, Vankelecom H. Protocol for establishing organoids from human ovarian cancer biopsies. STAR Protoc 2021;2. 10.1016/j.xpro.2021.100429.

[24] Hill SJ, Decker B, Roberts EA, Horowitz NS, Muto MG, Worley MJ, et al. Prediction of DNA repair inhibitor response in short-term patient-derived ovarian cancer organoids. Cancer Discov 2018;8:1404–21. 10.1158/2159-8290.CD-18-0474.

[25] Liston DR, Davis M. Clinically relevant concentrations of anticancer drugs: A guide for nonclinical studies. Clinical Cancer Research 2017;23:3489–98. 10.1158/1078-0432.CCR-16-3083.

[26] Gonzalez A, Nagel CI, Haight PJ. Targeted Therapies in Low-Grade Serous Ovarian Cancers. Curr Treat Options Oncol 2024;25:854–68. 10.1007/s11864-024-01205-4.

[27] Wang H, Wang L, Zhu X, He M, Zhong L, Lv Y, et al. Patient-derived organoids predict responses to chemotherapy and PARP inhibitors in advanced ovarian cancer. J Transl Med 2026;24:55. 10.1186/s12967-025-07112-y.

[28] Bruin MAC, Sonke GS, Beijnen JH, Huitema ADR. Pharmacokinetics and Pharmacodynamics of PARP Inhibitors in Oncology. Clin Pharmacokinet 2022;61:1649–75. 10.1007/s40262-022-01167-6.

[29] Tao M, Wu X. The role of patient-derived ovarian cancer organoids in the study of PARP inhibitors sensitivity and resistance: from genomic analysis to functional testing. Journal of Experimental & Clinical Cancer Research 2021;40:338. 10.1186/s13046-021-02139-7.

[30] Psilopatis I, Sykaras AG, Mandrakis G, Vrettou K, Theocharis S. Patient-Derived Organoids: The Beginning of a New Era in Ovarian Cancer Disease Modeling and Drug Sensitivity Testing. Biomedicines 2022;11:1. 10.3390/biomedicines11010001.

[31] Veneziani AC, Scott C, Wakefield MJ, Tinker A V., Lheureux S. Fighting resistance: post-PARP inhibitor treatment strategies in ovarian cancer. Ther Adv Med Oncol 2023;15. 10.1177/17588359231157644.

[32] Dong J, Ni J, Chen J, Wang X, Ye L, Xu X, et al. Genomic alteration discordance in the paired primary-recurrent ovarian cancers: based on the comprehensive genomic profiling (CGP) analysis. J Ovarian Res 2024;17:133. 10.1186/s13048-024-01455-8.

[33] Fehniger JE, Berger AA, Juckett L, Elvin J, Levine DA, Zajchowski DA. Comprehensive genomic sequencing of paired ovarian cancers reveals discordance in genes that determine clinical trial eligibility. Gynecol Oncol 2019;155:473–82. 10.1016/j.ygyno.2019.10.004.

[34] Glasgow MA, Argenta P, Abrahante JE, Shetty M, Talukdar S, Croonquist PA, et al. Biological insights into chemotherapy resistance in ovarian cancer. Int J Mol Sci 2019;20. 10.3390/ijms20092131.

[35] Newhouse R, Nelissen E, El-Shakankery KH, Rogozińska E, Bain E, Veiga S, et al. Pegylated liposomal doxorubicin for relapsed epithelial ovarian cancer. Cochrane Database Syst Rev 2023;7:CD006910. 10.1002/14651858.CD006910.pub3.

[36] Gorski JW, Zhang Z, McCorkle JR, DeJohn JM, Wang C, Miller RW, et al. Utilizing Patient-Derived Epithelial Ovarian Cancer Tumor Organoids to Predict Carboplatin Resistance. Biomedicines 2021;9:1021. 10.3390/biomedicines9081021.

[37] Moskalewicz M, Popova Y, Wiertlewska-Bielarz J. “From chemo to chemo”—the temporal paradox of chemotherapy. Supportive Care in Cancer 2021;29:3429–31. 10.1007/s00520-021-06039-6.

[38] Ortiz M, Wabel E, Mitchell K, Horibata S. Mechanisms of chemotherapy resistance in ovarian cancer. Cancer Drug Resist 2022;5:304–16. 10.20517/cdr.2021.147.

[39] Guffanti F, Mengoli I, Damia G. Current HRD assays in ovarian cancer: differences, pitfalls, limitations, and novel approaches. Front Oncol 2024;14. 10.3389/fonc.2024.1405361.

[40] Wang L, Wang X, Zhu X, Zhong L, Jiang Q, Wang Y, et al. Drug resistance in ovarian cancer: from mechanism to clinical trial. Mol Cancer 2024;23. 10.1186/s12943-024-01967-3.

