## Supplementary Figures for "Patient-Derived Organoids Functionally Stratify Epithelial Ovarian Cancer into Clinically Relevant Chemotherapy Response Phenotypes"

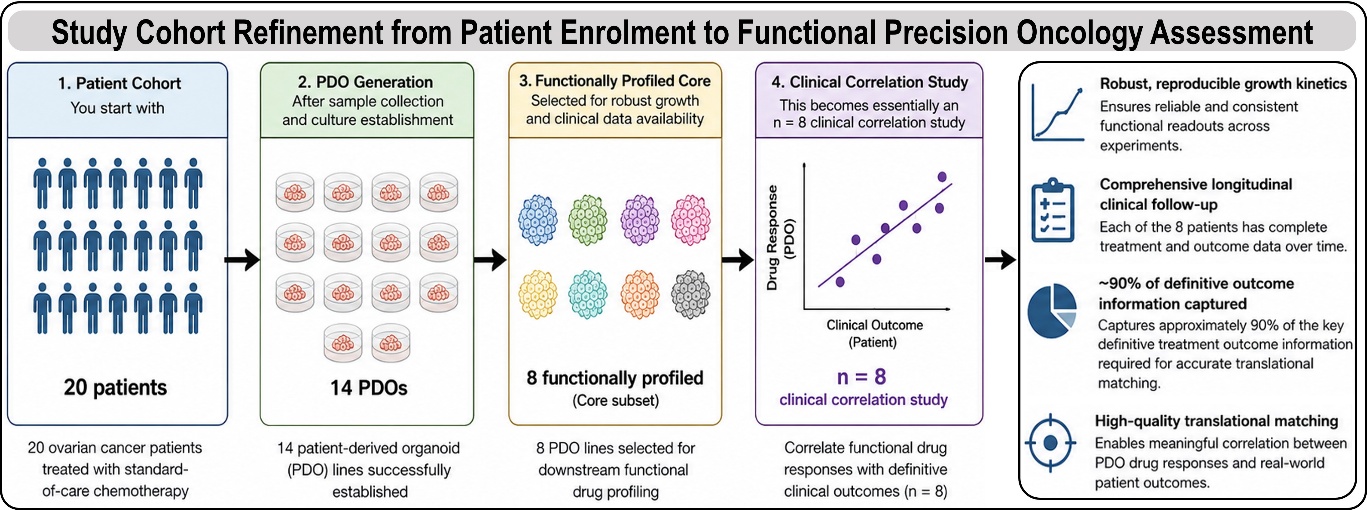

**Supplementary Fig. S1. Cohort selection and rationale for the effective clinical correlation cohort used for functional drug profiling.** A total of 20 epithelial ovarian cancer patients were enrolled for organoid establishment, from which 14 patient-derived organoid (PDO) lines were successfully generated and maintained as expandable cultures. Downstream functional drug profiling was subsequently restricted to a core subset of eight PDO lines exhibiting robust and reproducible growth kinetics together with comprehensive longitudinal clinical follow-up data. This curated subset captured approximately 90% of the definitive treatment outcome information required for translational matching, enabling reliable correlation between in vitro drug responses and patient clinical outcomes. Consequently, functional-clinical association analyses were effectively performed in an n = 8 cohort, prioritizing data completeness and biological reproducibility over cohort size.

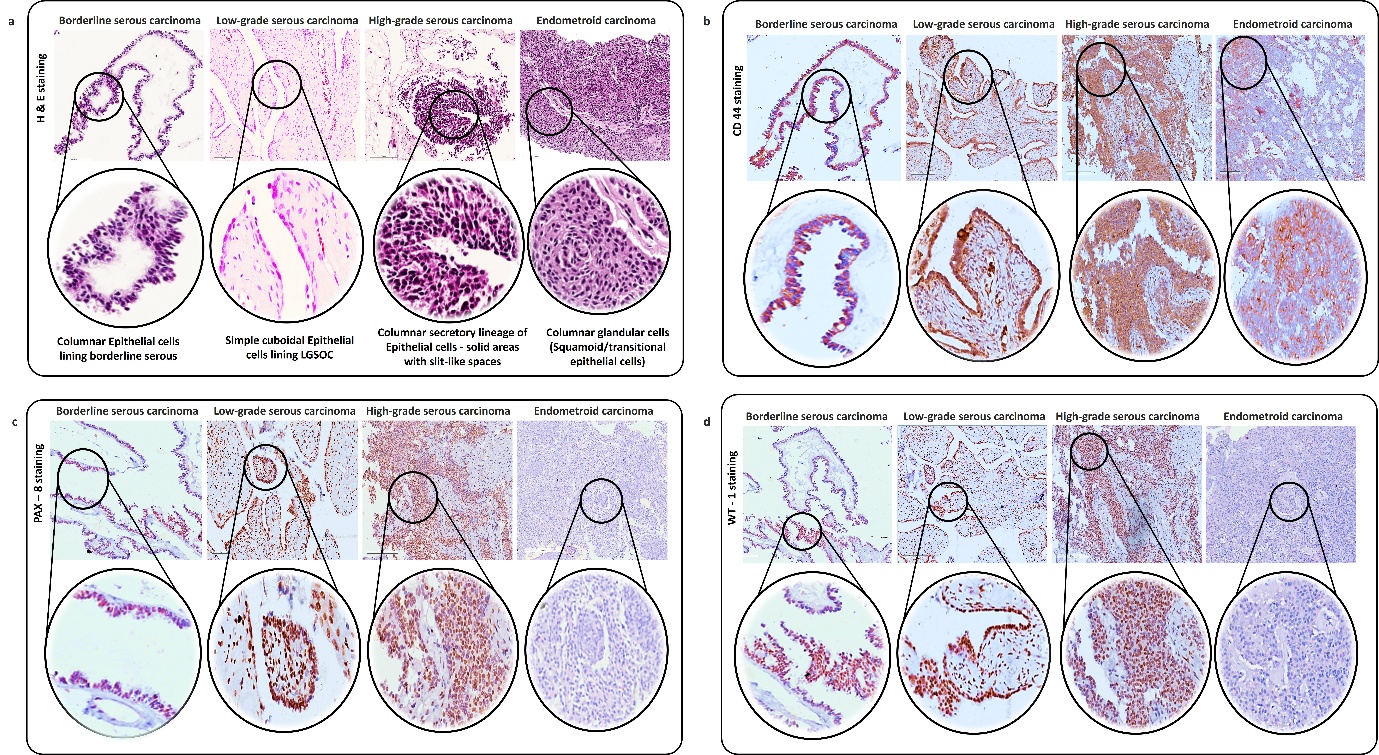

**Supplementary Fig. S2. Immunohistochemical profiling of primary ovarian tumour tissue.** Representative H&E and IHC staining of primary ovarian tumour sections for epithelial and tumour-associated markers (e.g., PAX8, WT1, CD44), confirming tumour identity and cellular heterogeneity. Patient tissue samples representing different histological types: POV-22 (Borderline serous carcinoma), POV-28 (Low-grade serous carcinoma), POV-13 (High-grade serous carcinoma), and POV-11 (Endometroid carcinoma) Sections were counterstained with haematoxylin. Scale bars, 50 µm.

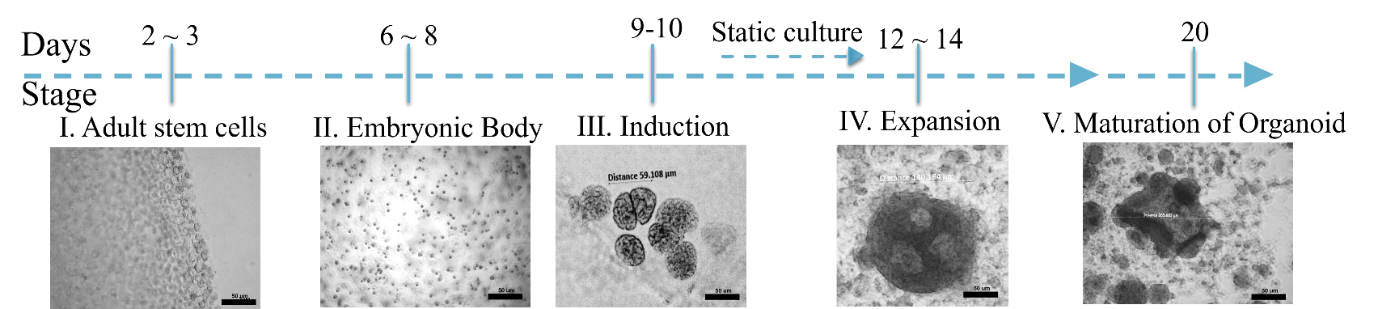

**Supplementary Fig. 3. Temporal progression of patient-derived organoid formation in 3D culture.** Representative bright-field images showing sequential stages of patient-derived organoid formation in 3D culture, from single-cell or small cluster seeding through early aggregation, spheroid formation, and maturation into structured organoids over time (timeline indicated). Scale bars, 50 µm.

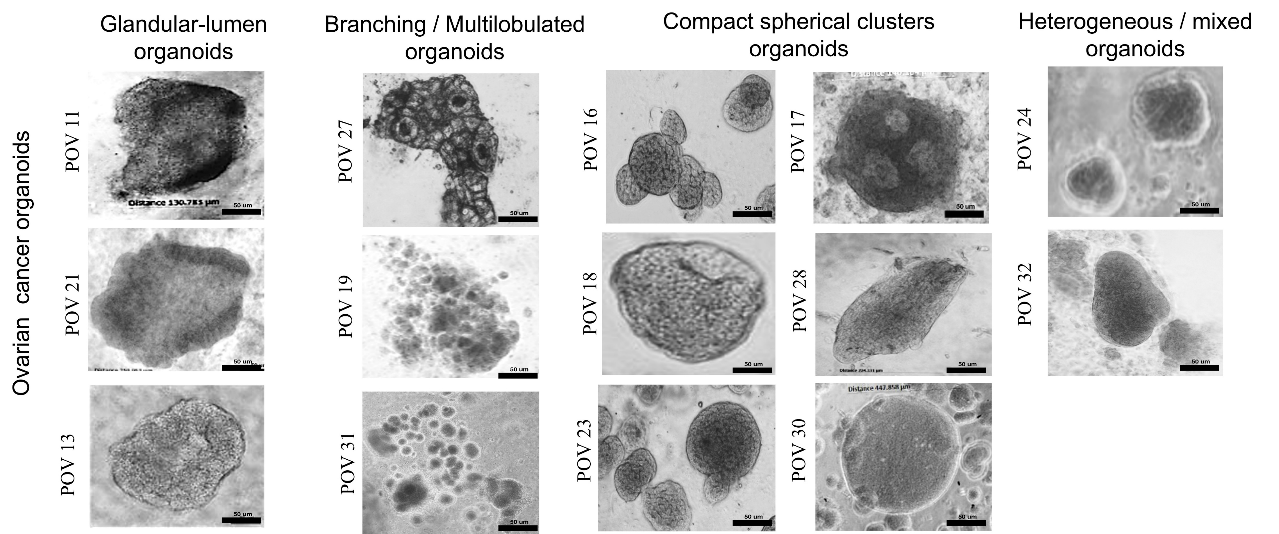

**Supplementary Fig. 4. Morphological heterogeneity of patient-derived ovarian cancer organoids.** Representative bright-field images of patient-derived ovarian cancer organoids demonstrating distinct morphological phenotypes (Sachs et al. 2018), including glandular-lumen structures, branching/multilobulated organoids, compact spherical clusters, and heterogeneous/mixed morphologies across different patient lines (POV IDs as indicated). Scale bars, 50 µm.

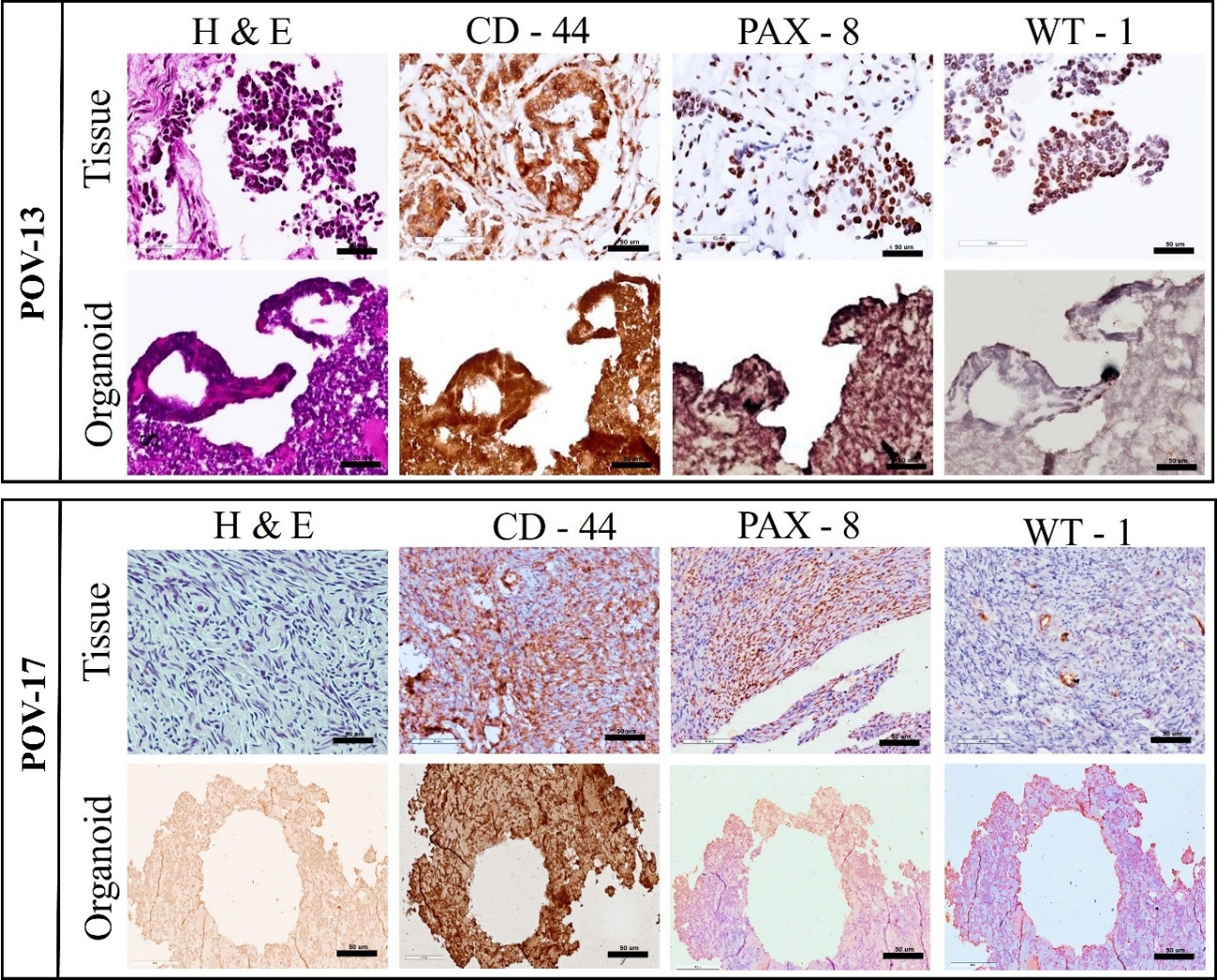

**Supplementary Fig. S5. Morphological features of patient-derived organoids by H&E staining.** Representative H&E staining and immunohistochemistry (IHC) of matched primary ovarian tumour tissue (top) and corresponding patient-derived organoids (bottom). PDOs recapitulate key histoarchitectural features observed in the parental tumours. IHC staining for CD44 (tumour/stemness-associated marker), PAX8 and WT1 (ovarian lineage markers) demonstrates concordant marker expression patterns between primary tissues and matched PDOs. Scale bars, 50 μm.

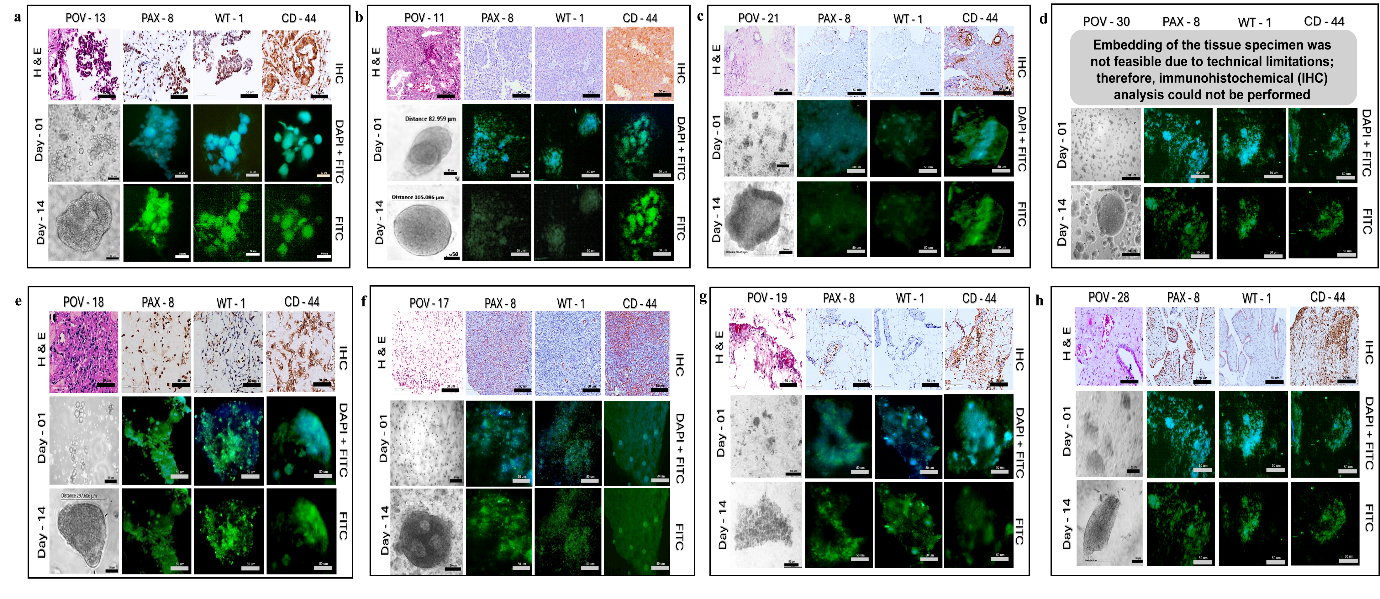

**Supplementary Fig. S6. Histological and immunophenotypic concordance between primary tumours and matched PDOs.** Representative haematoxylin and eosin (H&E) staining and immunohistochemistry (IHC) for lineage markers (PAX8, WT1) and the stem/progenitor marker CD44 in primary tumour sections, alongside whole-mount immunofluorescence (IF) of matched patient-derived organoids (PDOs) (POV IDs indicated). Corresponding bright-field and whole-mount immunofluorescence images of PDOs established from the same patients (POV-11, POV-13, POV-17, POV-18, POV-19, POV-21, POV-27, POV-28, and POV-30) are shown. PDOs were evaluated at early (Day 1) and established (Day 14) culture stages to assess preservation of tumour morphology and marker expression. DAPI (blue) was used as a nuclear counterstain. Representative bright-field images of PDO morphology are shown alongside immunofluorescence staining. For POV-30, embedding of the primary tumour specimen was not feasible due to technical limitations; consequently, immunohistochemical analysis could not be performed and only PDO images are presented. Scale bars, 50 μm.

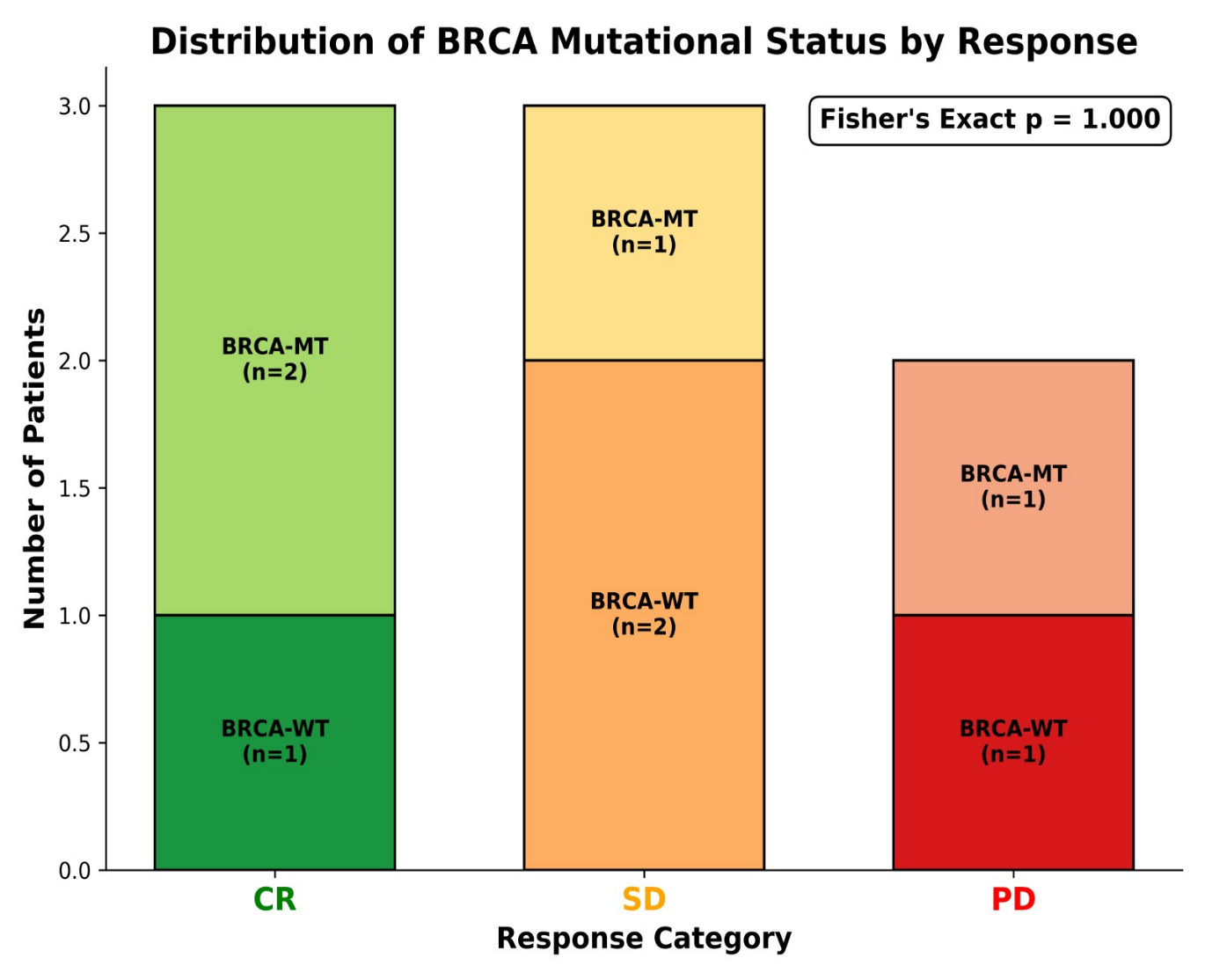

**Supplementary Fig. S7. Association of BRCA mutational status with clinical response in ovarian cancer.** Stacked bar plot showing the distribution of BRCA-mutant (BRCA-MT) and BRCA wild-type (BRCA-WT) cases across clinical response categories, including complete response (CR), stable disease (SD), and progressive disease (PD). Each bar represents the total number of patients within a response category and is subdivided according to BRCA mutational status, with the number of cases indicated within each segment. BRCA-mutant and BRCA wild-type tumors were observed across all response groups, with no significant association between BRCA status and clinical outcome (Fisher’s exact test, *p* = 1.000). These findings suggest that BRCA mutational status alone did not predict treatment response in this cohort, highlighting the need for complementary functional biomarkers to better capture therapeutic sensitivity and resistance.

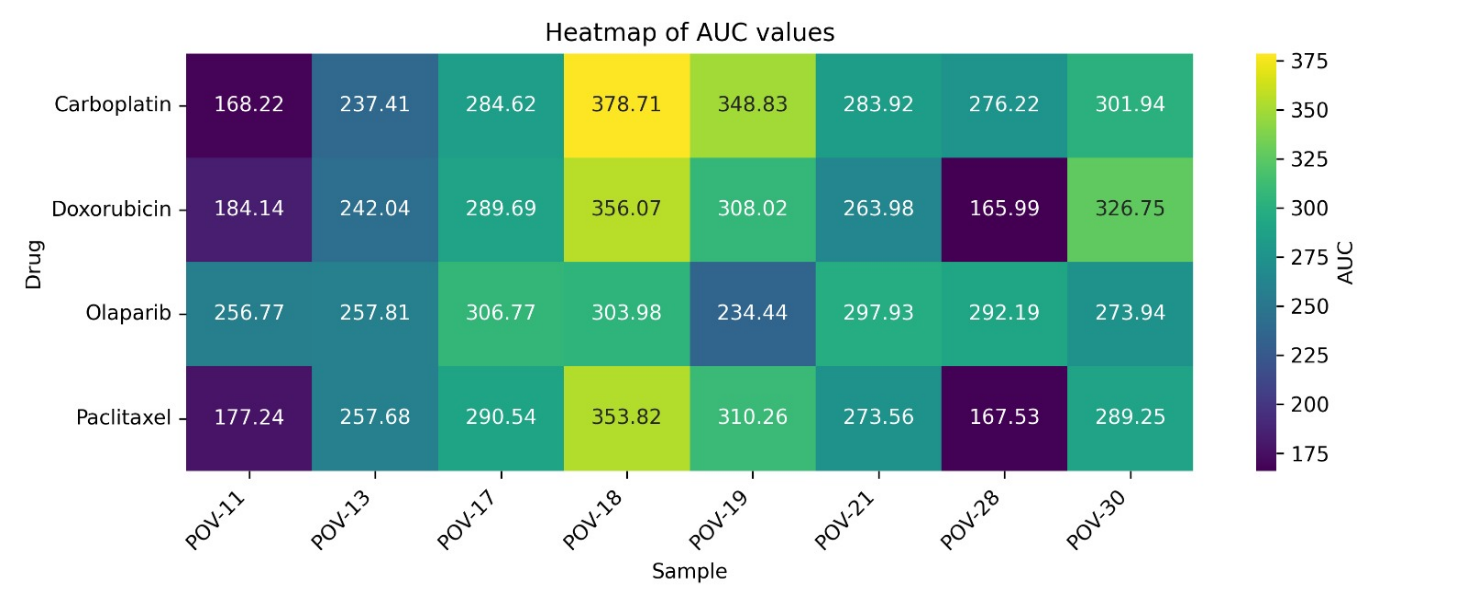

**Supplementary Fig. S8. Heatmap of normalized drug response (AUC) across patient-derived ovarian tumor organoids.** Heatmap showing normalized area under the curve (AUC) values (0–1) representing drug response across patient-derived ovarian tumor organoids (POV-11 to POV-30). Rows correspond to tested drugs (carboplatin, doxorubicin, olaparib, and paclitaxel), while columns represent individual organoid lines. Colour intensity reflects the relative AUC value, with lower values indicating greater drug sensitivity and higher values indicating relative resistance. The heatmap highlights heterogeneous drug response patterns across organoid models, demonstrating inter-patient variability in sensitivity to platinum-, taxane-, PARP inhibitor-, and anthracycline-based therapies.

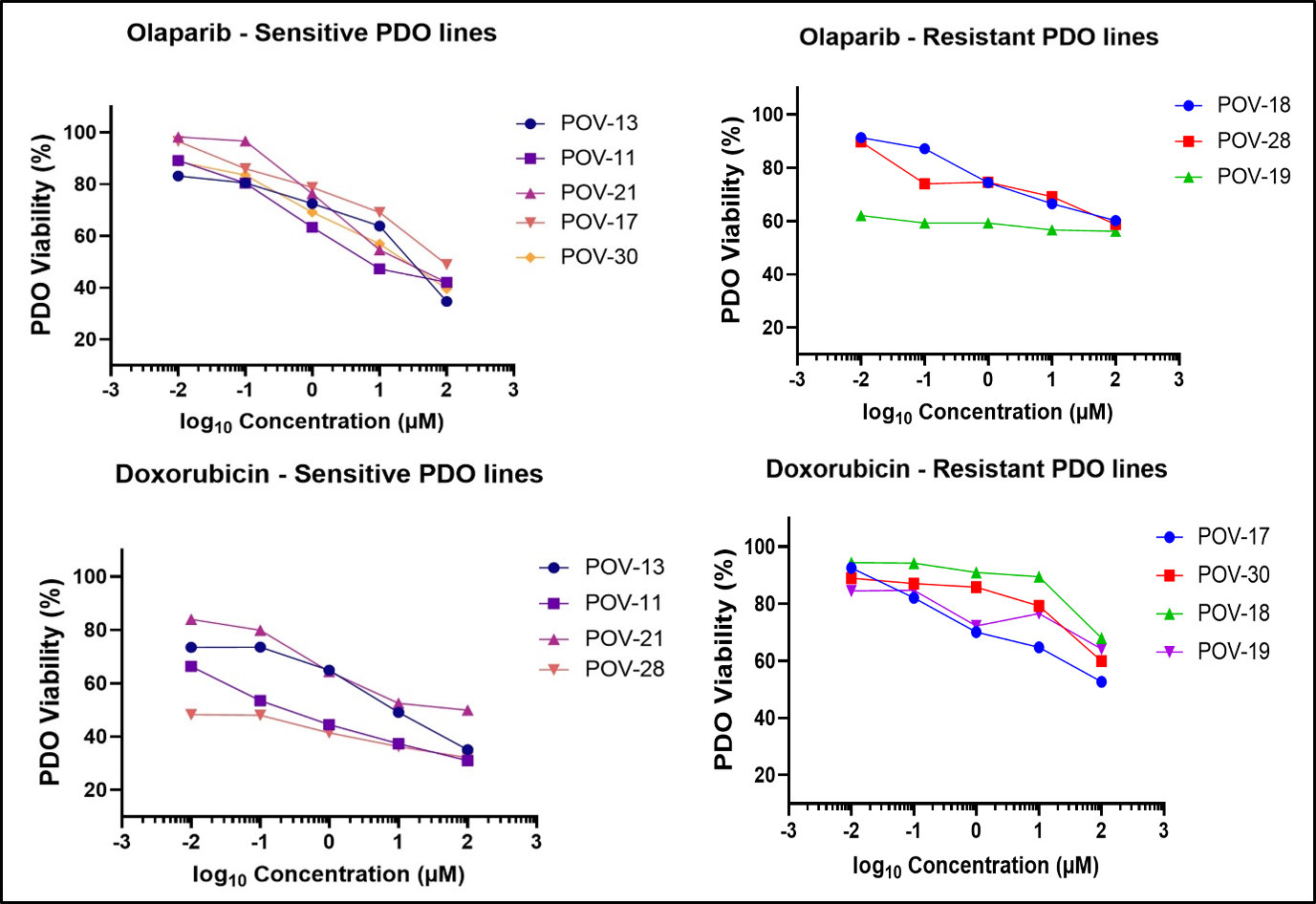

**Supplementary Fig. S9. Differential drug responses of ovarian cancer PDOs to olaparib and doxorubicin.** Dose-response curves depicting PDO viability (%) following exposure to increasing log_10_-transformed concentrations (µM) of olaparib and doxorubicin. Olaparib-sensitive PDOs (POV-13, POV-11, POV-21, POV-17, and POV-30) exhibited a dose-dependent decline in viability, whereas resistant PDOs (POV-18, POV-28, and POV-19) showed minimal response. Doxorubicin-sensitive PDOs (POV-13, POV-11, POV-21, and POV-28) demonstrated marked cytotoxicity, while resistant PDOs (POV-17, POV-30, POV-18, and POV-19) exhibited reduced sensitivity. These findings illustrate patient-specific differences in drug response and the functional heterogeneity of ovarian cancer PDOs.

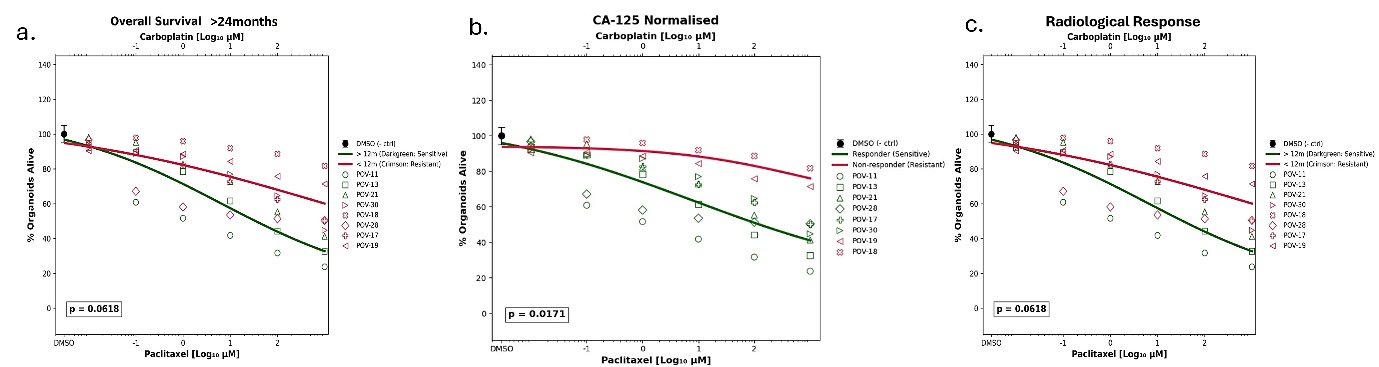

**Supplementary Fig. S10.** **Association of PDO carboplatin-paclitaxel response with clinical outcome measures.** Dose-response profiles of OC-PDOs treated with increasing concentrations of carboplatin-paclitaxel combinations and stratified according to clinical outcome parameters. The lower x-axis represents paclitaxel concentrations (log_10_ µM), while the upper x-axis represents the corresponding carboplatin concentrations (log_10_ µM). Organoid viability was normalised to DMSO-treated controls. Green symbols and fitted curves represent clinically sensitive PDOs, whereas red symbols and fitted curves represent clinically resistant PDOs. (A) Stratification according to overall survival (>18 months versus ≤18 months). (B) Stratification according to CA-125 normalization following treatment. (C) Stratification according to radiological response. Curves were generated using non-linear regression analysis of dose–response data. A significant separation between sensitive and resistant groups was observed for CA-125 normalisation (p = 0.0171), whereas associations with overall survival and radiological response did not reach statistical significance (p = 0.0618 for both comparisons).
