## Supplementary Results for "Patient-Derived Organoids Functionally Stratify Epithelial Ovarian Cancer into Clinically Relevant Chemotherapy Response Phenotypes"

*SR1. Tumour tissue validation and PDO formation*

Immunohistochemical assessment of primary tumour tissues confirmed adequate tumour cellularity prior to PDO establishment (**Supplementary Fig. S2**). Representative H&E and immunohistochemical staining demonstrated preservation of epithelial lineage markers and tumour-associated proteins across diverse histological subtypes.

Sequential bright-field imaging revealed a reproducible process of PDO formation, progressing from single cells or small clusters to mature organoid structures over 10 to 14 days in culture upon reaching 200-500 μm in diameter (Maenhoudt & Vankelecom, 2021) and remained amenable to cryopreservation and recovery, demonstrating long-term viability and stability (**Supplementary Fig. S3**).

*SR2. Morphological heterogeneity and histopathological concordance*

PDO cultures exhibited marked inter-patient morphological diversity, including glandular-lumen structures, branching and multilobulated organoids, compact spheroidal clusters, and mixed phenotypes (**Supplementary Fig. S4**).

Histological examination and immunophenotypic analyses demonstrated preservation of tumour architecture and expression of CD44, PAX8, and WT1 between parental tumours and matched PDOs. Whole-mount immunofluorescence further confirmed maintenance of lineage-specific marker expression during organoid expansion (**Supplementary Fig. S5 and Supplementary Fig. S6**).

*SR3. BRCA mutational status does not predict treatment response*

BRCA-mutant, BRCA-wild-type, and variant of uncertain significance cases were observed across complete response, stable disease, and progressive disease categories. No significant association was identified between BRCA mutational status and clinical outcome (Fisher’s exact test, p = 1.000; **Supplementary Fig. S7**), suggesting that genomic status alone incompletely captures therapeutic sensitivity.

*SR4. Reproducibility of functional drug profiling and multidrug response landscape*

Drug-response assays demonstrated high technical reproducibility with consistent AUC values between replicate experiments (**Supplementary 2 - Table S5**). Passage numbers at the time of screening are summarized in **Supplementary 2 - Table S6**.

Heatmap visualization of normalized AUC values highlighted substantial inter-patient variability in sensitivity to carboplatin, paclitaxel, olaparib, and doxorubicin, supporting the existence of heterogeneous therapeutic vulnerabilities among ovarian cancer PDOs (**Supplementary Fig. S8**).

*SR5. Differential responses to olaparib and doxorubicin*

Dose-response analyses revealed marked heterogeneity in olaparib sensitivity among BRCA-mutated PDOs. Olaparib-sensitive PDOs demonstrated progressive reductions in viability with increasing drug concentrations, whereas resistant models retained substantial viability across the tested dose range.

Similarly, doxorubicin-sensitive PDOs exhibited pronounced cytotoxicity, whereas resistant models maintained high viability, indicating persistence of tumour-intrinsic multidrug resistance mechanisms (**Supplementary Fig. S9**).

*SR6. Association between PDO carboplatin-paclitaxel sensitivity and clinical outcome measures*

Dose-response profiles stratified according to overall survival, CA-125 normalisation, and radiological response demonstrated significant separation between sensitive and resistant PDOs for CA-125 normalisation (p = 0.0171), whereas associations with overall survival and radiological response did not reach statistical significance (**Supplementary Fig. S10**).

These findings further support the relationship between PDO-derived functional responses and clinically relevant treatment outcomes.

*SR7. Individual patient treatment trajectories*

Detailed clinical histories, treatment timelines, maintenance strategies, and follow-up information for POV-11, POV-13, POV-17, POV-18, POV-19, POV-21, POV-28, and POV-30 are summarized in **Supplementary 2 - Table S1**.
