## Supplementary Methods for "Patient-Derived Organoids Functionally Stratify Epithelial Ovarian Cancer into Clinically Relevant Chemotherapy Response Phenotypes"

*S1. Patient cohort analytics and follow-up metrics*

Supplementary processing utilized clinical records and matched tissue specimens from the 20-patient OC cohort at Narayana Health City. The subset of eight PDO lines exhibited consistent linear expansion across early passages and was supported by longitudinal clinical data covering approximately 90% of the treatment period. Serum CA-125 kinetics were assessed using baseline measurements obtained before chemotherapy initiation and post-treatment values collected at a median follow-up of 4 weeks. The baseline clinicopathological attributes of the patient cohort are comprehensively cross-referenced in **Supplementary Table S1**.

*S2. Tissue dissociation and epithelial fraction enrichment*

Fresh surgical specimens (1-2mm^3^ fragments) were collected intraoperatively and transported to the laboratory in ice-cold Dulbecco’s Modified Eagle Medium (DMEM; Thermo Fisher Scientific, Cat. No. 11965092) supplemented with 1% penicillin-streptomycin. Tissues were washed three times in sterile PBS to clear red blood cells, mechanically minced with sterile cross-scalpels, and enzymatically digested in DMEM containing collagenase type IV (Sigma-Aldrich, Cat. No. C5138) at 37°C for 45-60 minutes under continuous, gentle orbital agitation. The resulting crude suspension was sequentially passed through a 70µm nylon cell strainer (Corning, Cat. No. 352350) to isolate single cells and small cell clusters from undigested stromal debris. Pellets were collected by centrifugation at 300xg for 5 minutes, treated briefly with RBC lysing buffer to eliminate residual erythrocytes, and resuspended in cold culture matrix.

*S3. Complete pdo culture formulation, passaging, and cryopreservation*

Enriched epithelial cells were resuspended in growth factor-reduced Extracellular Matrix (ECM) (Corning, Cat. No. 354277) at a 1:1 ratio and cast as 20µL droplets into pre-warmed 24-well plates. Following polymerization at 37°C for 15 minutes, domes were overlaid with 500µL of optimized ovarian cancer organoid medium. The basal medium consisted of Advanced DMEM/F12 (Thermo Fisher Scientific, Cat. No. 12491015) supplemented with 1x N2 (Cat. No. 17502048), 1xB27 minus vitamin A (Cat. No. 17504044), 1.25mM N-acetyl-L-cysteine, 10mM nicotinamide, 50ng/mL recombinant human EGF, 50ng/mL R-spondin-1, 100ng/mL Noggin, 500nM A83-01, 10µM SB203580, and 10µM Y-27632 dihydrochloride (added strictly during initial isolation and post-passaging transitions to mitigate anoikis). The exact working concentrations and vendor specifications for all medium components are provided in **Supplementary Table S2**.

Organoids were split 1:2 or 1:3 every 10-14 days. For passaging, matrix domes were disrupted with cold PBS, treated with TrypLE Express (Thermo Fisher Scientific, Cat. No. 12604013) at 37°C for 5 minutes, mechanically sheared, centrifuged, and re-seeded into fresh ECM domes. Long-term storage was achieved by dissociating organoids into single-cell suspensions, cryopreserving them in freezing medium, and storing them in liquid nitrogen vapor. Post-thaw viability was assessed by metabolic recovery.

*S4. Immunohistochemistry and immunofluorescence protocols*

PDO domes were harvested using Dispase, fixed in 4% paraformaldehyde (PFA) for 30 minutes, embedded in 2% low-melting-point agarose, and processed through graded alcohols into paraffin blocks. Sections (4-5µm) of both organoids and parental tissues were deparaffinized in xylene and rehydrated. Antigen retrieval was performed in a pressure cooker using Tris-EDTA buffer (pH 9.0) for 20 minutes. Endogenous peroxidase activity was quenched with 3% H_2_O_2_. Sections were blocked with 5% BSA and incubated overnight at 4°C with monoclonal antibodies against PAX8, WT1, and CD44. Signal development was executed via HRP-labelled secondary polymers and a DAB substrate kit, followed by counterstaining with hematoxylin.

For whole-mount immunofluorescence, intact organoids were fixed in situ in 4% PFA, permeabilized with 0.5% Triton X-100 for 1 hour, blocked with 5% BSA, and incubated with primary antibodies (1:400 dilution) for 16 hours at 4°C. Following three 20-minute washes in PBS containing 0.1% Tween-20 (PBS-T), organoids were incubated with FITC-conjugated secondary antibodies for 30 min. Subsequently, nuclei were counterstained with DAPI (1µg/mL). Slides were imaged using an epifluorescent microscope (Zeiss C, Axiocam and Zen lite 2012). Comprehensive antibody clones, dilution parameters, and full catalogue registries are structured in **Supplementary Table S3**.

*S5. Quantitative In Vitro Chemosensitivity Assays*

PDOs were collected, exposed to TrypLE to generate single-cell fractions, and filtered through a 40µm strainer. Cell density and viability were verified via trypan blue exclusion on an automated cell counter. Viable cells were suspended at a final concentration of 2,500-3,000 cells per well in a 50% Matrigel-medium matrix and seeded into 96-well optical-bottom plates.

Organoids were co-seeded with vehicle control or serial dilutions (0.01, 0.1, 1, 10, and 100 µM) of carboplatin, paclitaxel, doxorubicin hydrochloride, and olaparib. All assays were performed in technical triplicates. At 120 hours post-treatment, 50µL of activated XTT reaction mixture (prepared with phenazine methosulfate electron coupling reagent) was added per well. Plates were incubated at 37°C for 4 hours, and absorbance values were measured at 450nm (reference wavelength 630nm) on a Synergy H1 microplate reader (BioTek). Live/dead cell viability was assessed by staining cultures with a Calcein AM/Ethidium Homodimer-1 working solution for 15-20 min at room temperature, protected from light, followed by fluorescence imaging using an inverted microscope.

*S6. Dose-Response Modeling and Drug Sensitivity Metrics*

Raw absorbance entries were background-corrected against cell-free Matrigel blanks and normalized to the mean value of vehicle control wells (100% viability). Concentration-response parameters were computed using non-linear regression using a four-parameter logistic equation. IC_50_ values were derived from nonlinear dose-response curve fitting. Absolute AUC integrations across the standardized dosage domain (0.01-100µM) were executed in Python (v3.12.7) utilizing numerical integration formulas within the SciPy library. Clinical cutoff categorization utilized established human steady-state pharmacokinetic metrics (Cmax benchmarks: Carboplatin 145µM; Paclitaxel 4µM). Organoids displaying an IC_50_ value below these Cmax thresholds were classified as functionally sensitive.

*S7. Multi-modal translation and computational pipelines*

Clinical efficacy matching followed RECIST v1.1 classifications, mapping patient outcomes directly to corresponding ex vivo profiles. Baseline genomic profiles and mutational targets (e.g., *TP53*, *BRCA1/2*) are compiled in **Supplementary Table S4**.

All computational pipelines and downstream script generation were completed using Python (v3.12.7) utilizing the core analytical structures of pandas (v2.2.0), numpy (v1.26.4), and scikit-learn (v1.4.1), with data visualizations rendered through matplotlib (v3.8.3) and seaborn (v0.13.2). Comparative analysis evaluated paired CA-125 variations using two-tailed Wilcoxon signed-rank tests, and mutational significance across phenotypic clusters was parsed using exact contingency tests with significance set at p = 0.05.
